# Structure and mechanism of the tripartite ATP-independent periplasmic (TRAP) transporter

**DOI:** 10.1101/2022.02.13.480285

**Authors:** James S. Davies, Michael J. Currie, Rachel A. North, Joshua D. Wright, Mariafrancesca Scalise, Jack M. Copping, Daniela M. Remus, Ashutosh Gulati, Dustin R. Morado, Sam A. Jamieson, Gayan S. Abeysekera, Subramanian Ramaswamy, Rosmarie Friemann, Soichi Wakatsuki, Jane R. Allison, Cesare Indiveri, David Drew, Peter D. Mace, Renwick C.J. Dobson

## Abstract

In bacteria and archaea, tripartite ATP-independent periplasmic (TRAP) transporters uptake essential carboxylate- and sulfonate-containing nutrients into the cytoplasm. Unlike other secondary active transporters, TRAP transporters cannot receive their substrates directly, but do so indirectly *via* a secreted soluble substrate-binding protein. How a sodium-driven secondary active transporter is strictly coupled to a passenger-carrying substrate-binding domain is poorly understood. Here, we report the cryo-EM structure of the sialic acid TRAP transporter SiaQM from *Photobacterium profundum* at 2.97 Å resolution. SiaM has 12-TMs that come together to form a “transport” domain and a “scaffold” domain, with the transport domain consisting of helical hairpins as seen in the sodium-coupled elevator transporter VcINDY. Interestingly, the SiaQ protein forms intimate contacts with SiaM to extend the size of the scaffold domain, indicating TRAP transporters may operate as monomers, rather than the typically observed oligomers. We have identified the Na^+^ and sialic acid binding sites in SiaM and confirmed a strict dependence on the substrate-binding protein SiaP for uptake. We have determined the SiaP crystal structure that, together with co-evolution driven docking studies, provides a molecular basis for how sialic acid is delivered to the SiaQM transporter complex. We conclude that TRAP proteins are conceptually a marriage between an ABC importer and a secondary active transporter, which we describe herein as an ‘elevator-with-an-operator’ mechanism.

## Introduction

Transporter proteins play key roles in bacterial colonisation, pathogenesis and antimicrobial resistance^1–3^. Tripartite ATP-independent periplasmic (TRAP) transporters are a major class of secondary transporters found in bacteria and archaea—but, not in eukaryotes^4,5^. First reported over 20 years ago^6^, they use energetically favourable cation gradients to drive the import of specific carboxylate- and sulfonate-containing nutrients against their concentration-gradient, including C_4_-dicarboxylates, α-keto acids, aromatic substrates and amino acids^7^.

A functional TRAP system is made up of a soluble substrate-binding ‘P-subunit’, and a membrane-bound complex comprising a small ‘Q-subunit’ and a large ‘M-subunit’. For a small proportion of TRAP transporters, the Q- and M-subunits are fused into a single polypeptide^7–10^. TRAP transporters are different from almost all other secondary active transporters in that they can only accept substrates from the P-subunit^11^. Analogous to ABC importers, the P-subunit is secreted into the periplasm to capture host-derived substrates with high affinity and specificity. The substrate-loaded P-subunit is subsequently delivered to the membrane transporter QM^7–10^. The best characterised TRAP transporters are those for sialic acid, otherwise known as the SiaPQM system^12–14^. Sialic acids are a family of nine-carbon sugars of which the most common is *N-*acetylneuraminate (Neu5Ac)^12^. Sialic acid TRAP transporters have a demonstrated role in bacterial virulence^13–15^ and, as such, they represent an attractive class of proteins for the development of new antimicrobials against pathogenic bacteria that use them during infection. Despite their potential as antimicrobial targets and classification as evolutionary-divergent sodium-driven transporters, the molecular basis of how TRAP transporters work is poorly understood, with low sequence identity to transporters with known structures.

## Results

### The cryo-EM structure of SiaQM

The SiaQM transporter complex from *P. profundum*^16^ was selected for structural studies as it was found to be highly stable in detergent solution. To improve image alignment of the relatively small SiaQM complex for structural determination by cryo-EM^17^, synthetic nanobodies against SiaQM were generated using a yeast surface-display platform^18^. Promising nanobodies were converted into larger megabodies, which ultimately led to the selection of the megabody 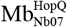 (see Methods). SiaQM is found to be a stable monomeric transporter complex (**Extended Data Fig. 1a-c**) and the size and shape is similar in either detergent (L-MNG) or amphipol (A8-35). The detergent purified SiaQM complex was forthwith exchanged into amphipol and combined with megabody to make the 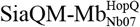 complex (**Extended Data Fig. 1d**). The sample preparation was next optimised for grid preparation, cryo-EM data acquisition and structural determination (see Methods). Final particles produced excellent 2D-classes with a 3D reconstruction yielding an overall resolution of 2.97 Å (FSC = 0.143 criterion), extending to 2.2 Å in some regions (**Extended Data Fig. 2, Extended Data Table 1**). Overall, 580 out of 597 residues of the SiaQM complex could be built, as expected from the high quality cryo-EM maps (**Extended Data Fig. 3)**.

The cryo-EM structure of the SiaQM complex reveals one copy of the Q-subunit and one copy of the M-subunit. The megabody has bound on the extracellular side of SiaQM, making contacts with both the Q- and M-subunits (**Fig. 1a**). The M-subunit is made up of 12 TM segments with an N_Out_C_Out_ topology. The M-subunit shows the highest structural similarity to the sodium-coupled dicarboxylate transporter VcINDY, which is a secondary active transporter operating by an elevator alternating-access mechanism^19^. Based on this structural comparison, we designate TMs 1-3 and TMs 7-9 as forming a “scaffold” domain, and TMs 4-6 and TMs 10-12 as forming a “transport” domain. The transport domain consists of two structural-inverted repeats. The first repeat contains a helical hairpin TM4a and TM4b originating from the cytoplasmic side (HP_in_), followed by the discontinuous helix TM5a-b and TM6. The second repeat has a helical hairpin from the periplasm side HP_out,_ and is made up from TM10a and TM10b, TM11a-b and TM12. The two symmetry-related repeats superimpose with an r.m.s.d. of 2.2 Å over 77 Cα positions. The scaffold domain is likewise made up from two structurally-inverted repeats, the first of which is formed by TM1, TM2 and TM3a-b, and the second by TM7, TM8 and TM9a (**Fig. 1b**). The scaffold structural-inverted repeats superimpose with an r.m.s.d. of 3.8 Å over 80 Cα positions and are connected to their neighbouring transport domains by lateral ‘arm helices’ that cradle the transport domain (**Fig. 1b**). The arm I helix appears to be locked in place by a salt-bridge formed between the conserved R75 in TM3c and D213 in TM7 of the scaffold I domain. R75 also forms cation-π interactions with Y254 in TM8 of the scaffold I domain (**Extended Data Fig. 4 and Extended Data Fig. 5**). The arm II helix 9b also appears to be stabilised by a cation-π interaction to TM9a in the scaffold II domain (**Extended Data Fig. 5**).

**Figure 1.**
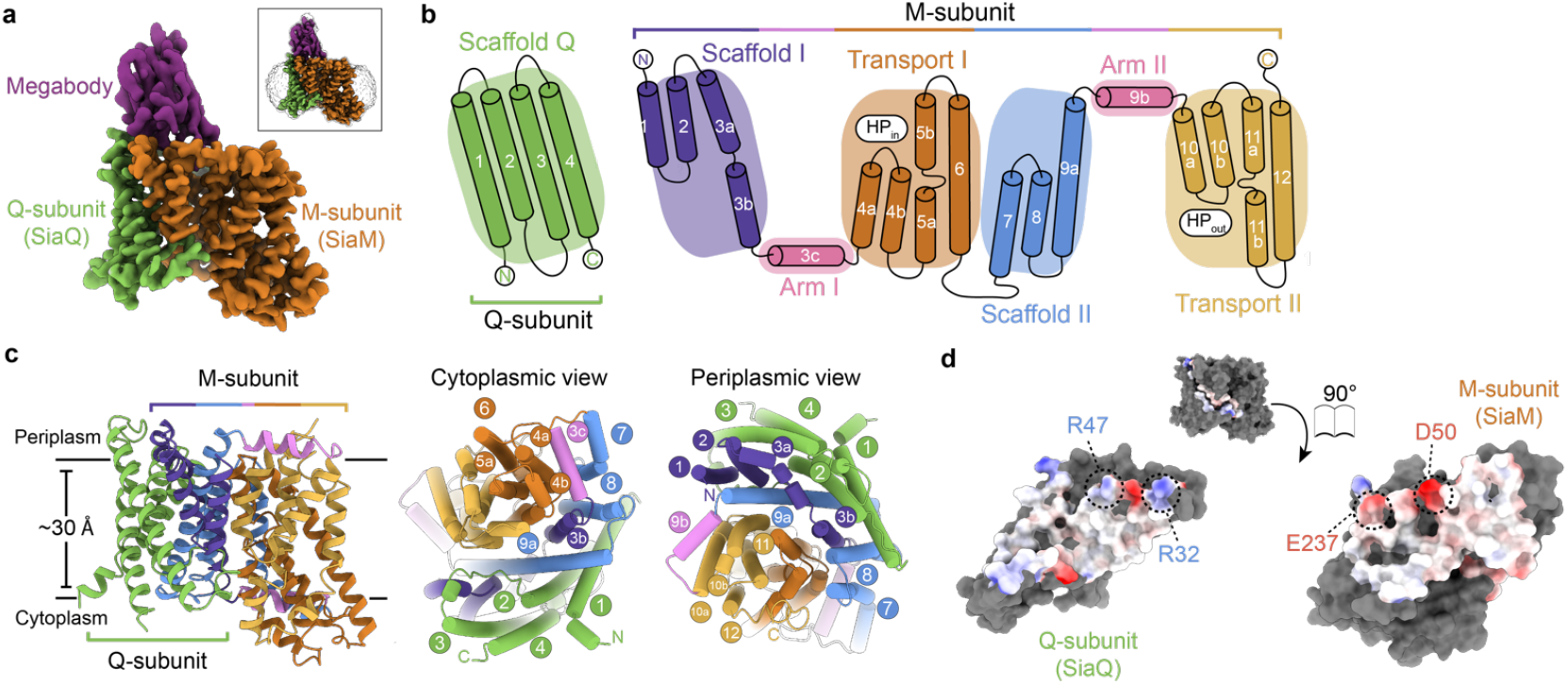
Structure of *Photobacterium profundum* SiaQM. **a**, Cryo-EM density for the 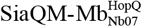 complex. The megabody is bound at the periplasmic face of SiaQM and makes contacts with both the Q- and M-subunits. *Inset* shows density and position of the amphipol belt. Cryo-EM density displayed at 7.5σ, as calculated by *ChimeraX*^37^. **b**, Cartoon depicting the topology of SiaQM. The overall topology of SiaQM can be arranged into a transport domain and a rigid scaffold domain. The inverted topology of the M-subunit is seen in this cartoon, where Scaffold I, Arm I and Transport I form one repeat, with the remainder forming the second repeat. Hairpin helices HP_in_ and HP_out_ are indicated. **c**, Structural organisation of SiaQM. SiaQM structure, showing the Q-subunit (green), and the M-subunit coloured by domain (scaffold domains: purple and blue; transport domains: orange and gold; arm helices: pink). Right: Views from the cytoplasm and periplasm, showing the scaffold domains (green, blue and purple) bracing the transport domains (orange and gold), which are cradled between the two arm helices (pink). **d**, Surface representation of the interface between SiaQ and SiaM. The surfaces of the contact residues at the interface are coloured by electrostatic potential, as calculated in *ChimeraX*^37^. Highlighted are conserved residues that form salt bridges between the subunits, R32:E237 and R47:D50. *Consurf*^38^ analyses show that R32 and E237 are fully conserved across the sequences sampled, highlighting the structural and functional significance of this pair.

The Q-subunit has an N_In_C_In_ topology as previously proposed^20,21^. It comprises three long helices (TM1, TM3 and TM4) and one shorter helix (TM2) with an extended cytoplasmic linker to TM3, and is positioned at an oblique angle (∼40º) relative to the membrane normal (**Fig. 1c**). The Q-subunit extensively interacts with the scaffold domain of the M-subunit, burying a total surface area of ∼2,400 Å^2^. SiaQ and SiaM interactions are dominated by van der Waals contacts, in addition to highly-conserved residues forming two salt bridges (**Fig. 1d**). The Q-subunit is enriched with tryptophan residues at the phospholipid-water interface (**Extended Data Fig. 4**), located near the termini of membrane spanning helices TM1, TM2 and TM4, which is a sequence feature seen in other TRAP transporters (**Extended Data Table 2**). In an elevator mechanism, the substrate is only translocated by the transport domain that moves against the scaffold domain, which is fixed due to oligomerisation^22^. Given the extensive interaction of the Q-subunit with only the scaffold domain of the M-subunit and the known role of tryptophan residues for anchoring helices in the membranes^23^, it seems likely the role of the Q-subunit is for extending the “scaffold” domain so that SiaQM can function as a monomer.

### SiaQM is electrogenic and SiaP dependent

To confirm the transport properties SiaPQM from *P. profundum* are equivalent to SiaPQM from *H. infleunzae*^24^, purified SiaQM was reconstituted into liposomes for transport assays. Uptake of ^3^[H]-Neu5Ac into SiaQM containing liposomes is indeed strictly dependent on both the presence of an inwardly directed Na^+^ gradient, and the presence of the soluble substrate-binding protein SiaP (**Fig. 2a-b)**. Net transport by SiaQM is electrogenic, as activity is enhanced when an inside negative membrane potential (ΔΨ) is imposed (**Fig. 2a**). Since Neu5Ac has a single negative charge at neutral pH, electrogenic transport means at least two Na^+^ ions are transported for every sialic acid molecule imported. Varying external Na^+^ at a fixed substrate concentration gives rise to a Hill coefficient of 2.7, implying possibly three Na^+^ ions are co-transported during each transport cycle (**Fig. 2c**).

**Figure 2.**
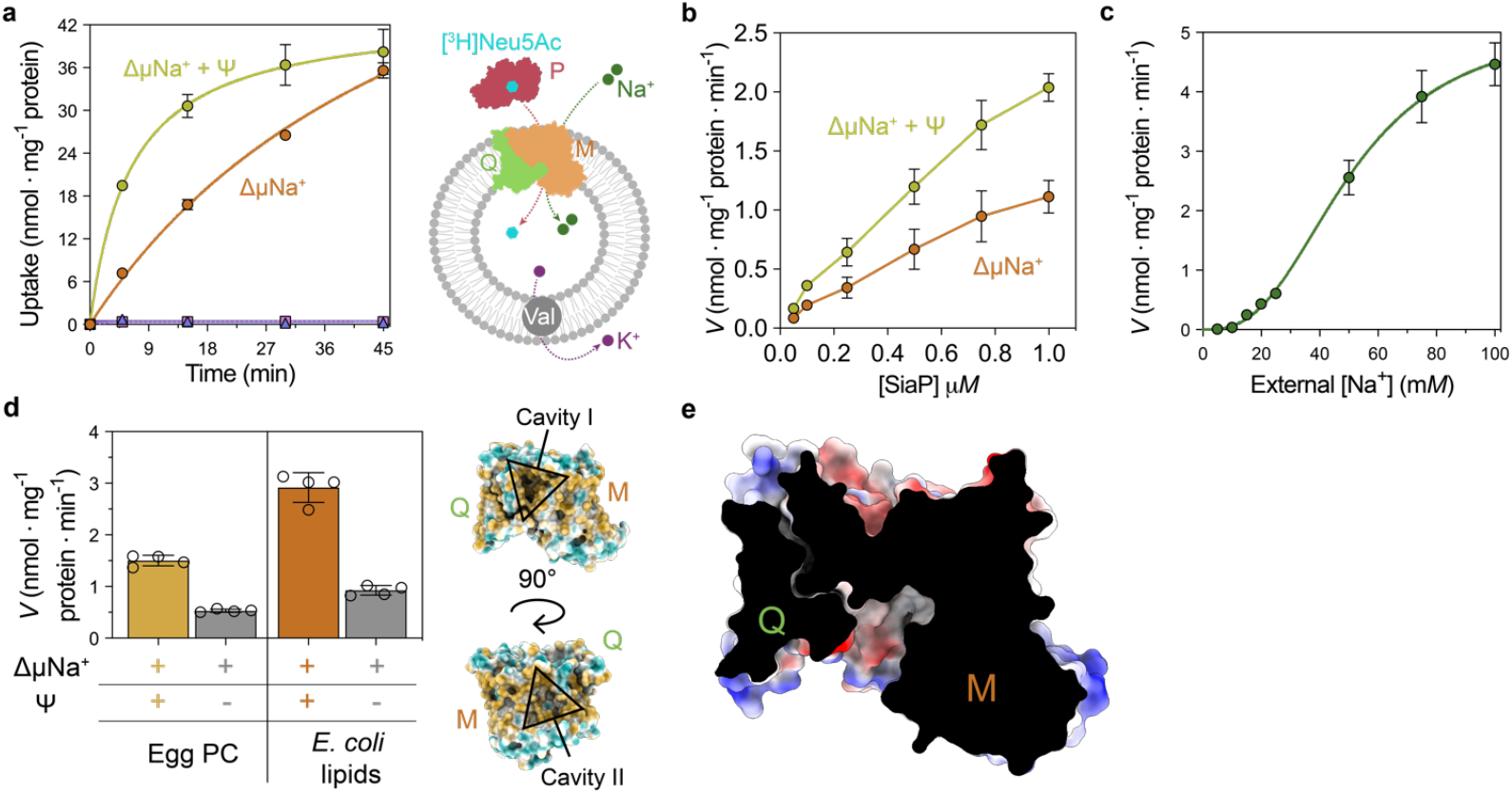
Proteoliposome transport assays of SiaPQM. **a**, Transport by SiaPQM is dependent on an inwardly directed Na^+^ gradient and net transport is electrogenic as activity is enhanced when an inside negative membrane potential (Ψ) is imposed. Curves show external [^3^H]-Neu5Ac uptake into proteoliposomes reconstituted with SiaQM, in the presence of SiaP. In 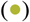 an inward sodium gradient (ΔμNa^+^) is present, with a membrane potential generated by valinomycin before measurement. In 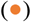 ethanol was added instead of valinomycin as a control. In 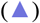 no ΔμNa^+^ was present, but a ΔΨ was imposed and in 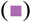 no ΔμNa^+^ was present and ethanol was added instead of valinomycin as a control. **b**, The dependence of Neu5Ac uptake into proteoliposomes based on SiaP. Transport was measured in the presence of varying concentrations of SiaP. The conditions of (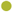 and 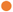) are the same as **a**. SiaP is required for transport but reduces the rate at a concentration of 1.0 μM. All data are reported as means ± s.e.m. from at least four independent experiments. **c**, Dependence of Neu5Ac uptake into proteoliposomes based on external Na^+^ 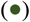. Transport was measured in the presence of varying concentrations of external Na^+^-gluconate and was fitted with the Hill equation giving a Hill coefficient of 2.7 (95% CI = 1.9–4.0), supporting Neu5Ac transport that is coupled to the translocation of two to three Na^+^ ions during each transport cycle. **d**, SiaQM transport activity is sensitive to the lipid environment. Transport was measured using proteoliposomes reconstituted with phosphatidylcholine from egg-yolk or *E. coli* total lipid extract. As a ΔΨ control, ethanol (grey bars) was used instead of valinomycin (orange bars). **e**, Surface cutaway of SiaQM. The structure is in an inward-facing conformation, which is the substrate release state, with the presence of a large solvent-accessible cavity on the cytoplasmic face of the complex.

In several elevator transporters, lipids have been shown to intercalate between the scaffold and transport domains, where the lipids may help to grease structural transitions^22,25,26^. Consistently, SiaQM transport activity is increased two-fold when proteoliposomes were prepared using *E. coli* phospholipid extract containing 70% of the non-bilayer forming lipid phosphatidylethanolamine, as compared with proteoliposomes prepared using phosphatidylcholine (**Fig. 2d**). Indeed, analysis of SiaQM’s surface shows a number of crevices and cavities that may bind lipids, with two conspicuous cavities positioned at the interface between the scaffold and transport domains, positioned adjacent to the arm helices (**Fig. 2d**).

### Substrate and sodium sites in SiaQM

The SiaQM structure is in an inward-facing conformation, as it has a large solvent-accessible cavity of 793 Å^3^ towards the cytoplasm (**Fig. 2e**). We identified two sodium ion sites in the SiaQM structure (**Extended Data Fig. 6a)**. The first sodium, Na1, is located between the loop of HP_in_ and the unwound region of TM5. The cryo-EM map shows weak map density that we attribute to a Na^+^ ion coordinated in a trigonal bipyramidal pattern by the five backbone carbonyls of S103, S106, G145, V148 and P150. We observe a second sodium site, Na2, located between the loop of HP_out_ and the loop between TM11a and TM11b. This second sodium site is less well-defined and the surrounding residues are not optimal for Na^+^ coordination. However, the backbone carbonyls of residues G325, G366, T369 and M372, and the sidechain hydroxyl of T369 are all positioned in close proximity and subtle re-configuration at this site would likely enable Na^+^ coordination. At each of the defined sodium sites is a highly-conserved twin proline motif, located at the peptide break of TM5a-b and TM11a-b, which was hypothesised to be required for sodium site formation^20^. Our structure confirms these prolines are juxtaposed to the Na^+^ ion-binding sites, and match those in the VcINDY structure, where the Na^+^ ions are positioned either side of the substrate^19,27^ (**Extended Data Fig. 6b**). Indeed, a surface topography analysis of the transport domain using CASTp^28^ identified a pocket with a solvent accessible volume of 79 Å^3^, which forms part of the large 793 Å^3^ vestibule (**Extended Data Fig. 6a**). Unlike other unrelated sodium-coupled sialic acid transporters that adopt the LeuT-fold^29,30^, the substrate binding-site is nearly absent of positively-charged residues for coordinating the negatively-charged sugar. Rather, the electrostatic surface potential shows a slight positive charge on one side of the cavity (above HP_in_), with a stronger negative charge opposite (below HP_out_) (**Extended Data Fig. 6a**). Based on the structure, it seems likely a conformational change in SiaM will be required to accept the substrate, which is consistent with the requirement of SiaP for transport. Indeed, despite sample preparation in the presence of 10 mM Neu5Ac for cryo-EM data collection, there is no detectable density for the substrate.

### Coupling of SiaP to SiaQM

Uptake of sialic acid is dependent on the concentration of the substrate-binding protein SiaP (**Fig. 2b**). The binding affinity (*K*_D_) for the interaction between SiaQM and SiaP is determined to be ∼400 μM in the presence of sialic acid by microscale thermophoresis (see Methods, **Extended Data Fig. 1e**). In analytical ultracentrifugation experiments, however, no SiaP binding to SiaQM could be detected in amphipol (Methods, **Extended Data Fig. 1f**). It is possible that the SiaP and SiaQM interaction is too transient and/or SiaM is preferentially in an inward-facing conformation in amphipol, which is unable to interact with SiaP. Indeed, despite extensive trials in both detergent and amphipol, we have been unable to obtain a cryo-EM structure of a complex between SiaP and SiaQM.

Using an alternative approach, we determined the crystal structure of SiaP bound to Neu5Ac in a closed state at 1.04 Å resolution, as a crystal structure of substrate-loaded SiaP enables us to model the SiaPQM complex (**Extended Data Fig. 7, Extended Data Table 3**). The algorithms *RaptorX*^31^, *Gremlin*^32^ and *AlphaFold*^33^ were found to predict similar contacts, where the P-subunit interacts with both the Q- and M-subunits (**Fig. 3a**). The periplasmic surface of SiaQM shows a bowl-like shape with a large area where SiaP docks. The bowl is lopsided, with the lip at the side of the Q-subunit higher than that of the M-subunit. Predicted contacts involve mainly surface residues of the scaffold domain and can be used to orient the P-subunit with respect to the Q- and M-subunits (**Fig. 3a**).

**Figure 3.**
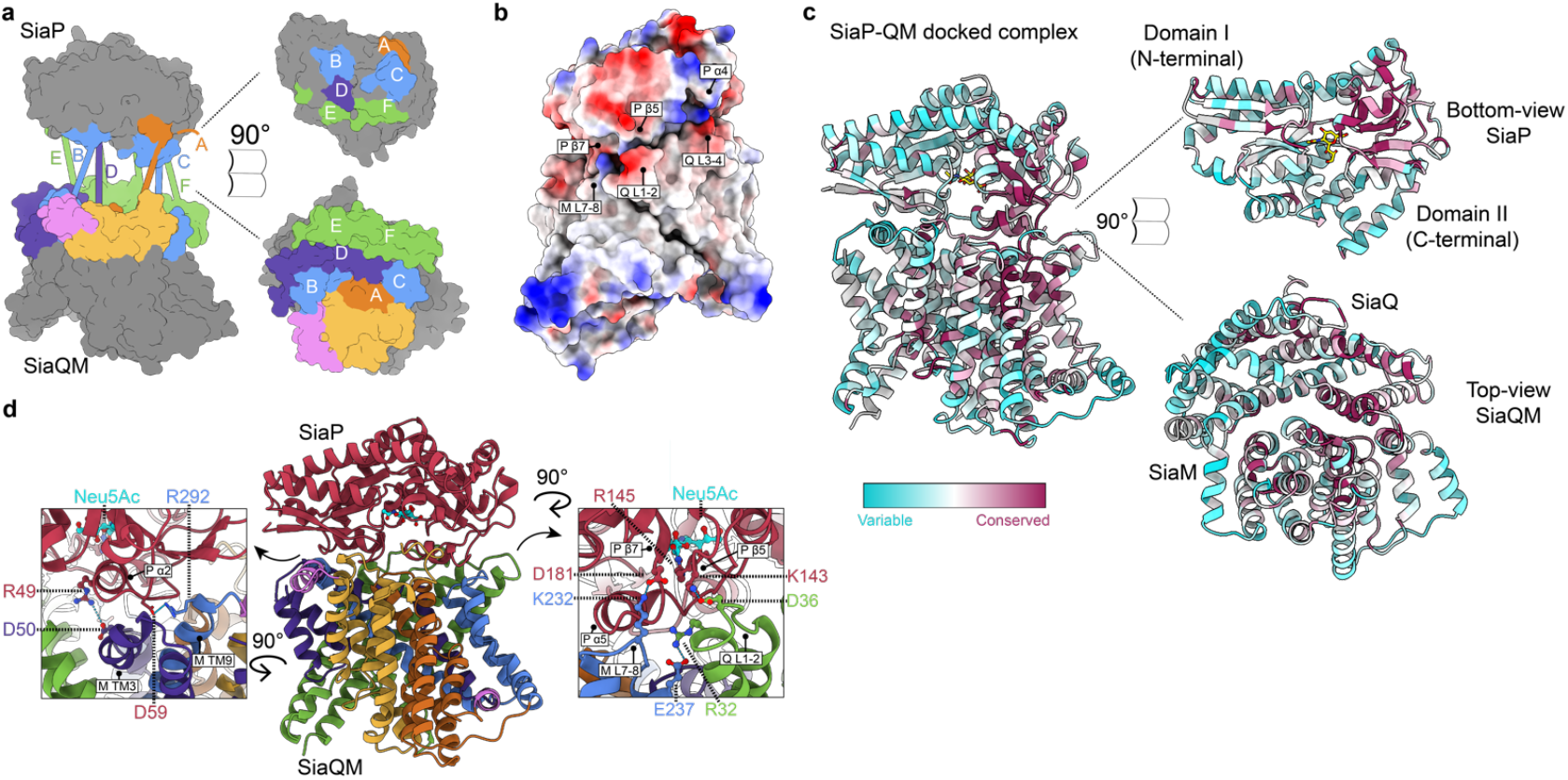
Predicted full TRAP complex. **a**, Interacting regions of SiaQM and SiaP, as determined using the algorithms *RaptorX*^31^, *Gremlin*^32^ and *AlphaFold*^33^ (**Extended Data Table 4-5**). Surface representation of the SiaQM and SiaP structures with patches coloured and lettered on each surface to indicate the binding mode. The proposed binding surface on SiaQM largely involves the surface of the scaffold (green, purple and blue). TM3a aligns well with α2 of the P-subunit (B blue and D purple). The loop between TM7 and TM8 aligns well with α5 of the P-subunit (C blue). The two periplasmic loops of the Q-subunit are also predicted to interact with the P-subunit (E and F, green). The coloured patches contain residue pairs with high co-evolution signal between the proteins, as represented by the solid bars on the left. The orange and pink patches represent the Transport I and Arm II components as coloured in **Fig. 1b. b**, Surface representation of the complex, coloured by electrostatic potential. Highlighted are regions of each structure we identify as interaction hotspots. **c**, Residue conservation mapped onto the SiaP and SiaQM structures. Cartoon representation of SiaP and SiaQM, coloured according to *Consurf* score^38^. The structure is opened (right) to show the conservation at the interacting surfaces, with a large cluster of conserved residues on the surface of the C-terminal domain of SiaP (specifically around η4-α5 helix, β6, β7 and η5). On the corresponding surface of SiaQM there is also a cluster of conserved residues (around Q L1-2, M L5b-6, M L7-8). **d**, Complex of the SiaQM cryo-EM structure and SiaP crystal structure based on the binding mode predicted by *AlphaFold2*^39^. An interaction hotspot (inset, right) shows a number of titratable residues at the interface. The interaction of the loop between TM1 and TM2 of the Q-subunit (Q L1-2), with the β5 strand of the P-subunit is of interest, as β5 houses the conserved arginine (R145) that forms a salt bridge with the ligand, Neu5Ac. The surface-accessible, highly conserved lysine near β5 (K143), two residues upstream of R145, is well-positioned to interact with the Q-subunit, and may therefore be involved in both recognition and allosteric modulation of the P-subunit. The invariant aspartic acid group (D181) of the P subunit also interacts with a lysine (K232) of the M-subunit. A similar interaction is also seen on the opposite side of the complex (inset, left) where R292 of TM9a of the scaffold is well-positioned to interact with D59 of SiaP. Also depicted is a hotspot involving D50 at the unwound region of TM3 in the scaffold portion of the M-subunit (see **Fig. 1b** for topology), which is well-positioned to interact with R49 of the P-subunit (D purple in **Fig. 3a**).

In the SiaP structure, Neu5Ac is situated in a deep cleft bound by multiple residues, including R145, which is highly conserved in TRAP transporter substrate-binding proteins^7,8^ (**Extended Data Fig. 7a-c**). Two calliper-like helices of SiaP, α2 at the N-terminal lobe and α5 at the C-terminal lobe, interact with the scaffold portion of the M-subunit (**Extended Data Fig. 7a**). Specifically, TM3a aligns well with α2 of the P-subunit, as does the loop between TM7 and TM8 with α5 of the P-subunit (**Fig. 3a**). The two periplasmic loops of the Q-subunit are also predicted to interact with the P-subunit, and analysis of the surface charge shows complementarity at this contact position, particularly at the loop between TM3 and TM4 (**Fig. 3a-b**). The predicted interacting surfaces show high levels of sequence conservation (**Fig. 3c**). Contacts from the transport domains are localised to the short helix TM5b in the Transport I domain, and the loop connecting TM5b to TM6, which has predicted contacts with SiaP at its short 3_10_ helix (η5) spanning residues F195 to E197.

Next, we used the docked model to predict how Neu5Ac is delivered to SiaQM for transport. Importantly, in the model we observe regions of surface and charge complementarity, sequence conservation and a high co-evolution signal, consistent with the docking and allosteric opening of SiaP with SiaQM. Clear hotspots involving potential salt-bridges are formed between the scaffold domain and the N- and C-terminal lobes of SiaP, likely contributing to both recognition and allosteric modulation of the P-subunit (**Fig. 3d**). Of note is an interaction between K232 on the loop between TM7 and TM8 (L7-8) of the M-subunit and D181 of SiaP immediately prior to β7 (**Fig. 3b and d**, right). Adjacent to this interaction, the residue D36 in the loop between TM1 and TM2 of the Q-subunit, interacts with the conserved K143 at β5 of SiaP (**Fig. 3b and d**, right). This strand in SiaP extends from the protein surface to the Neu5Ac binding site, with K143 located close to R145 directly coordinating Neu5Ac. Thus, SiaP binding to SiaQM is likely to pull on the strand next to the D36-K143 salt-bridge to release the substrate.

### An elevator-with-an-operator mechanism

The M-subunit shares a conserved fold with DASS members VcINDY, LaINDY and NaCT, as well as the bacterial AbgT-type transporter^34^ (**Extended Data Fig. 8**). These transporters undergo elevator-structural transitions, which involve a vertical translation and a rigid-body rotation of the transport domain against the scaffold^35^. Molecular dynamics simulations of LaINDY show that elevator-like motions induce the bending of flexible loops connected to the arm helices of the hairpin helices in the transport domains^35^. Due to a shared fold and the rigidity of the arms in our SiaQM structure, we suggest TRAP proteins undergo similar conformational changes. Likewise, similar to the DASS members, there are very few interactions between the transport and scaffold domains, further supporting our proposal that the M-subunit moves as a rigid body independently against the scaffold domain. Here, it seems that the extra scaffolding provided by the Q-subunit enables the transporter to uniquely function as a monomer. It is possible that the larger scaffold may have arisen to accommodate the interaction with SiaP, which could have become hindered upon an oligomeric assembly of SiaM.

Although the transporter component of TRAP transporters shares no sequence or structural similarities with ABC transporters, the mechanism of substrate translocation shares elements with Type I ABC importers^36^. Essentially, the P-subunit is analogous to the substrate-binding protein of ABC importers, which bind substrates with very high affinity and specificity. The substrate-binding protein then passes the substrate to the transporter, which otherwise has no, or poor, affinity for the substrate on its own. The consistent observation that substrate translocation requires the P-subunit is strong evidence that the substrate-binding protein modulates the SiaQM structure. Critically, the predicted binding interaction mode between the P-subunit and SiaQM is over the scaffolding domain. Almost certainly, the binding of SiaP triggers a conformational change in SiaQM that will enable the transport domain to accept the substrate, although it is unclear whether this is a local, gating rearrangement and/or a conformational switch from an inward-to an outward-facing conformation. Subsequently, substrate-free SiaP releases its interaction with SiaQM, which enables the protein to re-capture another substrate molecule. We propose that the TRAP transport cycle can be described as an ‘elevator-with-an-operator’ mechanism (**Fig. 4**). Supposedly, the TRAP system has evolved to enable the capture of scarce nutrients, but with high affinities (very slow off-rates) that would otherwise be incompatible with transporters that require efficient turnover by binding substrates weakly, in the μM-to-mM range. In conclusion, our work highlights how the transport cycle of a small molecule transporter can be controlled by the donation of the substrate from a secreted protein, greatly expanding our general view of secondary active transport.

**Figure 4.**
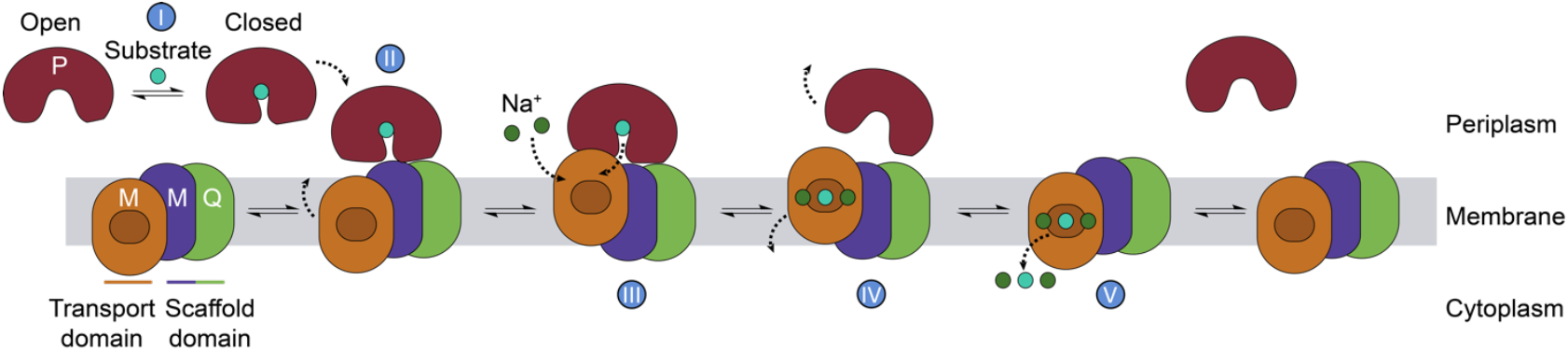
The TRAP ‘elevator-with-an-operator’ mechanism. (I) The P-subunit (maroon) binds the substrate (cyan) with high affinity and undergoes a conformational change from the open to closed state. (II) The closed P-subunit then docks to the QM subunits (orange, purple and green). (III) We propose that docking induces a conformational change in the transporter to a state where Na^+^ ions (green) and the substrate can bind with greater affinity. This change is coupled to the allosteric modulation of the P-subunit to the open conformation, releasing the substrate to the transporter. (IV) The open state P-subunit, which presumably has lower affinity for the transporter, diffuses away, allowing the transporter to move to an inward-facing state (V), with the substrate and coupling ions then released into the cytoplasm. We note it is possible the conformational change induced in the transporter (II) may be either a local gating rearrangement, or a global elevator type motion. Regardless, we suggest that the P-subunit is the ‘operator’ of the elevator, as transport without the P-subunit is negligible, as seen in **Fig. 2b**.

## Acknowledgements

This research was undertaken in part using the MX2 beamline at the Australian Synchrotron, part of the Australian Nuclear Science and Technology Organisation (ANSTO), and made use of the Australian Cancer Research Foundation (ACRF) detector as well as the SAXS beamline at the Australian Synchrotron, part of ANSTO. R.C.J.D., R.A.N., J.R.A., S.W., J.S.D. and M.J.C acknowledge funding support from the Marsden Fund, managed by Royal Society Te Apārangi (contract UOC1506) and the Biomolecular Interaction Centre (UC). R.C.J.D., J.S.D. and M.J.C. also acknowledge the following for funding support, in part: 1) the Ministry of Business, Innovation and Employment Smart Ideas grant (contract UOCX1706); 2) the Maurice Wilkins Centre flexible research grant; and 3) the Australian Institute of Nuclear Science and Engineering (AINSE Ltd) and ANSTO for a Postgraduate Research Award. R.A.N. acknowledges the Canterbury Medical Research Fund (contract CMRF 08). J.R.A. acknowledges the Rutherford Discovery Fellowship, managed by the Royal Society Te Apārangi (contract 15-MAU-001/15-UOA-008). J.C. is supported by a University of Auckland Doctoral Scholarship. The authors acknowledge the facilities, and scientific and technical assistance from flow cytometry staff at Otago Micro and Nanoscale Imaging (OMNI), at the University of Otago. We thank Prof. Borries Demeler (University of Lethbridge, Canada) for help with AUC experiments. R.F. acknowledges the Swedish Governmental Agency for Innovation Systems (2017-00180), and the Centre for Antibiotic Resistance Research (CARe) at the University of Gothenburg. C.I. acknowledges the MIUR (Ministry of Education, University and Research) Italy for the support through the “SI.F.I.PA.CRO.DE. – Sviluppo e industrializzazione farmaci innovativi per terapia molecolare personalizzata PA.CRO.DE.” (PON ARS01_00568). D.D acknowledges funding from the Knut and Alice Wallenberg Foundation. Cryo-EM data were collected at the Cryo-EM Swedish National Facility funded by the Knut and Alice Wallenberg Foundation, the Family Erling Persson and Kempe Foundations, SciLifeLab, Stockholm University and Umeå University.

## Author contributions

Project designed by R.A.N. and R.C.J.D. Research funding was obtained by R.C.J.D, R.A.N., P.D.M., J.R.A. and S.W. Protein expression and purification were performed by J.S.D., M.J.C., R.A.N., J.W., D.M.R. and G.A. Grid optimisation was carried out by R.A.N. and Cryo-EM data collection was performed by R.A.N and D.R.M. Cryo-EM data processing, map refinement and 3D reconstruction were performed by R.A.N. and A.G. Model building and structure building was carried out by J.S.D., with refinement, analysis and interpretation by J.S.D. and M.J.C. Nanobody screening and selection was performed by S.A.J. and P.D.M. Nanobody production and megabody design for Cryo-EM was carried out by J.S.D. AUC and MST experiments were conducted and analysed by M.J.C. Proteoliposome assays were performed and analysed by M.S. and C.I. X-ray crystallography experiments, data processing, and structure determination were performed by J.S.D. and R.A.N. Refinement was performed by J.S.D and J.W. Bioinformatic analysis was performed by J.W. and J.S.D. Structural modelling, docking and interpretation of the complex was performed by J.S.D and J.C. Overall experimental development, analysis of data and interpretation of the results was overseen by R.A.N., S.R., R.F., S.W., J.R.A., C.I., D.D., P.D.M., and R.C.J.D. The manuscript was prepared by J.S.D., M.J.C., R.A.N., D.D., and R.C.J.D, with contributions from all authors.

## Extended data for

## Methods

### Protein expression and purification

The sequence encoding the genetically non-fused *Photobacterium profundum* SiaQM (**Extended Data Table 6**) was cloned into pBAD-HisA (GeneArt) using XhoI and EcoRI restriction sites so that a hexahistidine affinity tag was fused at its 5’ end. The sequence retained the native intergenic region (5’-GGATTTTTC-3’) between the Q- and the M-subunits. This plasmid was transformed into *Escherichia coli* TOP10 cells (Invitrogen). Cells were grown at 37 °C to log phase at an OD_600_ of 1.7-1.9 in Terrific-Broth (TB). Recombinant protein expression was induced with 0.2% arabinose for 3 h. Cells were collected and homogenised in phosphate buffered saline (PBS) pH 7.4, 0.5 mg mL^-1^ lysozyme, 0.1 mM PMSF and lysed by ultrasonication at 70% amplitude in 0.5 sec on, 0.5 sec off cycles for 30 min, using a Hielscher UP200S Ultrasonic Processor. Cell debris was removed by two centrifugation steps at 16,000 *g* and 4 °C for 30 min. Membranes were harvested by ultracentrifugation at 240,000 *g* in a 50.2 Ti rotor (Beckman Coulter) at 4 °C for 2 h, and solubilised in PBS pH 7.4, 6% glycerol, 5 mM DTT, 0.1 mM PMSF and 2% w/v lauryl maltose neopentyl glycol (L-MNG, Anatrace) for 2 h. Insoluble material was removed by ultracentrifugation at 160,000 *g* in a 50.2 Ti rotor (Beckman Coulter) at 4 °C for 1 h. SiaQM was purified by Ni^2+^-NTA affinity using a 5 mL HisTrap HP column equilibrated in 50 mM Tris-HCl pH 8.0, 150 mM NaCl and 0.002% (w/v) L-MNG. Solubilised material was loaded onto the column and washed with 20 column volumes (CVs) of equilibration buffer, and bound protein was eluted in 10 CVs of equilibration buffer supplemented with 500 mM imidazole. Eluted protein was concentrated for size-exclusion chromatography (SEC) using 100 kDa molecular weight cut-off (MWCO) spin concentrators (Pall). SEC was carried out using a HiLoad 16/60 Superdex 200 size-exclusion column (Cytiva) in buffer comprised of 50 mM Tris-HCl pH 8.0, 150 mM NaCl and 0.002% w/v L-MNG. SEC fractions containing SiaQM were pooled and concentrated. Protein was either used in subsequent experiments, or flash-frozen in liquid nitrogen and stored at -80 °C for future use.

The sequence encoding SiaP without the native signal peptide (residues 1−22, from SignalP^40^) (**Extended Data Table 6**) was cloned into pET30ΔSE and transformed into *E. coli* BL21(DE3). Cells were grown at 37 °C to log phase at an OD_600_ of 0.6−0.8 in Luria-Broth (LB). Recombinant protein expression was induced with IPTG (1 mM) for 16 h at 26 °C. Cells were collected and lysed by ultrasonication at 70% amplitude in 0.5 sec on, 0.5 sec off cycles for 10 min. Cell debris was removed by centrifugation at 20,000 *g* and 4 °C for 20 min. SiaP was purified by anion-exchange chromatography using a Q 16/10 anion exchange column (Cytiva) equilibrated in 50 mM Tris-HCl pH 8.0. The column was washed with 5 CV equilibration buffer and bound protein was eluted with a gradient to 50 mM Tris-HCl pH 8.0, 1 M NaCl across 10 CV. Ammonium sulfate was added to the eluted SiaP to a final concentration of 1 M. SiaP was further purified by hydrophobic-interaction chromatography and bound to a Phenyl FF 16/10 column (Cytiva) equilibrated in 50 mM Tris-HCl pH 8.0 and 1 M ammonium sulfate. The column was washed with 5 CV equilibration buffer and bound protein was eluted with a gradient to 50 mM Tris-HCl pH 8.0, 150 mM NaCl across 10 CV. Eluted protein was concentrated for SEC using 10 kDa MWCO spin concentrators (Pall) and loaded onto a HiLoad 16/60 Superdex 200 size-exclusion column (Cytiva) equilibrated with 50 mM Tris-HCl pH 8.0, 150 mM NaCl and 1 mM *N*-acetylneuraminic acid (Carbosynth). SEC fractions containing SiaP were pooled and concentrated. Protein was either used in subsequent experiments, or flash-frozen in liquid nitrogen and stored at -80 °C for future use.

### Reconstitution of SiaQM from *Photobacterium profundum* in proteoliposomes

Purified SiaQM was reconstituted using a batch-wise detergent removal procedure as previously described^29,30^. In brief, 50 μg of SiaQM was mixed with 120 μL 10% C_12_E_8_, 100 μL of 10% egg yolk phospholipids (w/v), in the form of sonicated liposomes as previously described^41^ (except where differently indicated in the figure legend), 50 mM of K^+^-gluconate, 20 mM HEPES/Tris pH 7.0 in a final volume of 700 μL. The reconstitution mixture was incubated with 0.5 g Amberlite XAD-4 resin under rotatory stirring (1200 rev/min) at 25 °C for 40 min^41^.

### Transport measurements and transport assay

After reconstitution, 600 μL of proteoliposomes were loaded onto a Sephadex G-75 column (0.7 cm diameter × 15 cm height) pre-equilibrated with 20 mM HEPES/Tris pH 7.0 with 100 mM sucrose to balance the internal osmolarity. Then, valinomycin (0.75 μg/mg phospholipid), prepared in ethanol, was added to the eluted proteoliposomes to generate a K^+^ diffusion potential, as previously described^29^. After 10 sec of incubation with valinomycin, transport was started by adding 5 μM [^3^H]-Neu5Ac to 100 μL proteoliposomes in the presence of 50 mM Na^+^-gluconate and 0.5 μM of SiaP. For kinetic measurement, the initial transport rate was measured by stopping the reaction after 15 min, *i*.*e*., within the initial linear range of [^3^H]-Neu5Ac uptake into the proteoliposomes as determined by time course experiments. The transport assay was terminated by loading each proteoliposome sample (100 μL) on a Sephadex G-75 column (0.6 cm diameter × 8 cm height) to remove the external radioactivity. Proteoliposomes were eluted with 1 mL 50 mM NaCl and collected in 4 mL of scintillation mixture, vortexed and counted. The radioactivity taken up in controls performed with empty liposomes, *i*.*e*., liposomes without incorporated protein, was negligible with respect to the data obtained with proteoliposomes, *i*.*e*., liposomes with incorporated proteins. Data analysis was performed by Grafit software (version 5.0.13) using the Hill plot for kinetics determination and the first-rate order equation for time course analysis. All measurements are presented as means ± s.e.m. from independent experiments as specified in the figure legend.

### Crystallography, data collection and analysis

Purified SiaP in SEC buffer was concentrated to 50 mg mL^−1^ using a 10 kDa MWCO spin concentrator and crystallised using the sitting-drop vapour-diffusion method at 20 °C. The best diffracting crystals were grown in the E1 condition from the Shotgun screen (2.0 M ammonium sulfate, 0.1 M Bis-Tris pH 5.0) (Molecular Dimensions). Crystals were cryo-protected by soaking in reservoir solution supplemented with liquid glycerol and ethylene glycol.

Crystallographic data were collected on the MX2 beamline at the Australian Synchrotron at a wavelength of 0.95372 Å, using an Eiger 16M detector (ACRF ANSTO). Data were scaled and processed using *XDS*^42^ and *CCP4*^43^. Phases were obtained by molecular replacement using Phaser^44^ with 2xwk as the search model. Model building was performed using *COOT*^45^, and refinement performed with *Phenix*^46^. Atomic displacement parameters (ADPs) were refined anisotropically. PDB-redo was used to optimise the model. The final refined model had 98.49% favoured and zero outliers in the Ramachandran plot. The structure was deposited into the PDB with identification code 7t3e.

### Nanobody screening and megabody design

Nanobodies were selected using methods modified from McMahon *et al*.^18^. In short, SiaQM protein was purified and modified with amine reactive FITC- and Alexafluor647-labels (A647; ThermoScientific). Labelling was verified using spectrophotometry and equated to approximately 1.8 and 2.0 fluorescent label per protein molecule, respectively. Nanobodies were selected in buffer comprised of 20 mM HEPES pH 7.6, 150 mM sodium chloride, 0.1% (w/v) bovine serum albumin, 5 mM maltose, 0.002% L-MNG and 2 mM sialic acid. To negatively select non-specific clones, the naïve nanobody library was incubated with 400 μL anti-A647 magnetic beads (Miltenyi Biotec), immobilised in an LD column, and the flowthrough incubated with 1 μM A647 labelled SiaQM. SiaQM-binders were enriched by adding 400 μL of anti-A647 magnetic beads (or anti-FITC beads as appropriate), and captured in LS columns (Miltenyi Biotec), then released into Yglc4.5-Trp media for recovery. After one day of recovery, yeast were transferred to -Trp glucose media, and after 24 hours nanobody expression was induced with galactose for 24 hours prior to selection. The selection procedure encompassed two passes of three selection types—successive rounds of magnetic selection with anti-A647, then anti-FITC protein capture, and a third selection by fluorescence assisted cell sorting (FACS) using A647-labelled protein. Selection was progressively more stringent over these rounds, with protein concentrations of, 1 μM, 500, 200, 60, 30, 10 nM. After the final round of FACS, a dilution series of the recovered cells were plated on YPD agar. Single colonies were grown in 96-well plates and amplified for Sanger sequencing using a modified primer pair (for-GAAGGTGTTCAATTGGACAAGAGAGAAGCTGAC; rev-GCGTAATCTGGAACATCGTATGGGTAGGATCC). Sequences were aligned using *Geneious*^47^. Nanobodies were selected for follow-up based on enrichment at the sequence level and confirmation of labelled SiaQM binding to individual yeast clones assessed by flow cytometry. The successful sequence of Nb07 comprised twelve of the ninety-six (12.5%) sequenced clones from the final sequenced plate.

The megabody 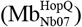 was constructed using a reported protocol^17^, as follows: residues 1-13 of the nanobody β-strand A (residues 1-13), followed by the C-terminal domain of *Helicobacter pylori* HopQ (Uniprot ID: B5Z8H1) (residues 227−446), which was directly fused to the N-terminal domain of HopQ (residues 53−221) followed by the remainder of the nanobody (residues 14−122) (**Extended Data Table 6**). The sequence encoding the megabody 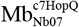 was synthesised by Genscript, and cloned into the expression vector pET-22B(+), encoding an N-terminal *pelB* signal peptide for periplasmic targeting and a C-terminal hexahistidine tag. Protein overexpression and purification were carried out using established methods^17^.

### Amphipol exchange

SEC purified SiaQM was incubated with amphipol A8-35 (Anatrace) at a 1:5 w/w ratio for 2 hours before addition of 100 mg mL^-1^ Bio-Beads SM-2 resin (Bio-Rad) and then incubated overnight at 4 °C with gentle agitation to remove detergent. After exchange, SEC was performed in 50 mM Tris pH 8.0 and 150 mM NaCl buffer to remove free amphipol and assess protein monodispersity.

### Megabody complex formation

SiaQM in amphipol was complexed with the megabody 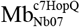 at a 1:1.5 molar ratio. SEC using a 24 mL Superdex 200 increase 10/300 GL was used to further purify the 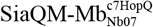 complex, in 50 mM Tris-HCl pH 8.0, 150 mM NaCl. Fractions containing the 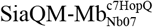 complex (**Extended Data Fig. 9**) were concentrated, flash-frozen in liquid nitrogen and stored at -80 °C for cryo-EM experiments.

### Assessment of complex formation

Sedimentation velocity analytical ultracentrifugation was used to assess complex formation with three samples. SiaQM (3.5 μM) in amphipol, 5.25 μM 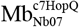, and both combined (1:1.5 molar ratio). The size-exclusion buffer was 50 mM Tris-HCl pH 8, 150 mM NaCl.

### Single-particle cryo-EM vitrification and data acquisition

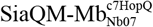 was concentrated to 2.9 mg mL^-1^ in the presence of 10 mM sialic acid, and 3 μL of sample applied to a freshly glow-discharged (Gatan Solarus) Quantifoil R2/1 cu300 mesh grid (Electron-microscopy Sciences). Grids were blotted using a Vitrobot Mark IV (Thermofisher Scientific) for 3.5 sec, at 4 °C, and with 100% humidity before vitrification in liquid ethane. Cryo-EM datasets were collected on a Titan Krios G3i electron microscope equipped with a K3 detector and BioQuantum imaging filter (Gatan), operated at 300 kV and in counting mode. Movie stacks were collected at a nominal magnification of 130,000× and a 0.6645 Å pixel size, with a dose rate of 16.57 e^−^/pixel/sec. Each movie was a result of 1.9 sec exposures with a total accumulated dose of 71.3 e^−^/Å^2^, which were fractionated into 40 frames. The EPU software package (Thermo Fisher Scientific) was used for automated data collection and all movies were recorded with a defocus range of −2.0 to −0.4 μm in 0.2 μm increments. An energy filter slit width of 20 eV and a 50 μm C2 condenser aperture were used during imaging, while the objective aperture was not inserted. In total, 10,960 micrographs were recorded and the statistics of cryo-EM data acquisition are summarised in **Extended Data Table 1**.

### Single-particle cryo-EM data processing and map refinement

Unless otherwise stated, all cryo-EM data processing was performed using *CryoSPARC* v.3.2.0^48^ (**Extended Data Fig. 1**). Movie frames were aligned using patch motion correction with a *B*-factor of 500, then contrast transfer function (CTF) estimations were made using the patch CTF estimation tool. Initially, 229,273 particles were picked from 300 micrographs using the Blob Picker tool and extracted. These particles were 2D classified into 50 classes, and 5 of these 2D classes were selected and used as a template for automated particle picking using the Template Picker tool, where 7,173,112 particles were picked from 10,960 micrographs. Particles were inspected with the Inspect Picks tool using an NCC Score Threshold of 0.12, and a Local Power range of 22,000 to 54,000. A total of 5,609,037 particles were extracted with a box size of 300 pixels and Fourier cropped to 200 pixels. The extracted particles were then sorted using iterative rounds of 2D classification, where the best 32 classes showing some structural details were selected, retaining 1,067,809 particles. The particles were subjected to *Ab initio* reconstruction separated into three classes. The best 3D reconstruction contained 624,033 particles and was used as a reference model for an initial round of non-uniform refinement, allowing a 3.29 Å model to be reconstructed. Particles were re-extracted with a final box size of 300 pixels and Fourier cropped to 240 pixels, giving a final pixel size of 0.83 Å. Iterative rounds of local refinement followed, using a mask that included SiaQM and nanobody, and excluding the surrounding amphipol, using an initial low pass resolution of 10 Å, searching over a range of 1° in orientations and 1 Å shifts.

### Structural model building and analysis

The atomic model of 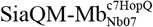 was built *de novo* from the globally sharpened 2.97 Å map using the *phenix* Map to Model tool^49^. Segments were manually joined using *coot* (version 0.9.5 EL), and the structure was refined using *phenix* real space refine using secondary structure, rotamer and Ramachandran restraints. The *Namdinator* tool^50^ was used between rounds of manual model building to optimise geometry and reduce clashes. Models were validated using Molprobity within *phenix* ^48^. The SiaQM:SiaP complex was predicted using *AlphaFold*^33^, with experimentally determined structures then aligned to this complex. *Rosetta fast relax*^51,52^ was then used to improve the sidechain packing of the complex. The structure was deposited into the PDB with identification code 7qha and the EMDB with the identification code EMD-13968.

### Microscale thermophoresis

SiaQM in amphipol was labelled at a ratio of 0.8 dye / 1 protein following the protocol of the RED-NHS 2^nd^ generation labelling kit (NanoTemper). Labelled SiaQM was held constant at 20 nM and SiaP was titrated at 16 concentrations starting at 2,900 μM and serially diluting 2-fold to 90 nM. The buffer contained 50 mM Tris pH 8, 150 mM NaCl and 10 mM Neu5Ac. Capillaries were loaded into a Monolith NT.115 (NanoTemper) and measured at 60% LED with a red detector, medium laser power and 22 °C. The experiment was performed in duplicate.

### Analytical ultracentrifugation

Sedimentation velocity experiments were performed using an XL-I analytical ultracentrifuge (Beckman Coulter). Reference solution (400 μL) and sample solutions (380 or 400 μL) were loaded into cells with double sector 12-mm Epon centrepieces and sapphire windows. The samples were run in an An-60 Ti rotor at 42,000 rpm and 20 °C until all species had completely sedimented. Both absorbance (280 or 450 nm) and interference optical systems were used to monitor sedimentation. Data analysis was performed with *Ultrascan 3* (version 4.0)^53^, *Sedfit*^54^, and *GUSSI*^55^.

## Extended Data Figures

**Extended Data Figure 1.**
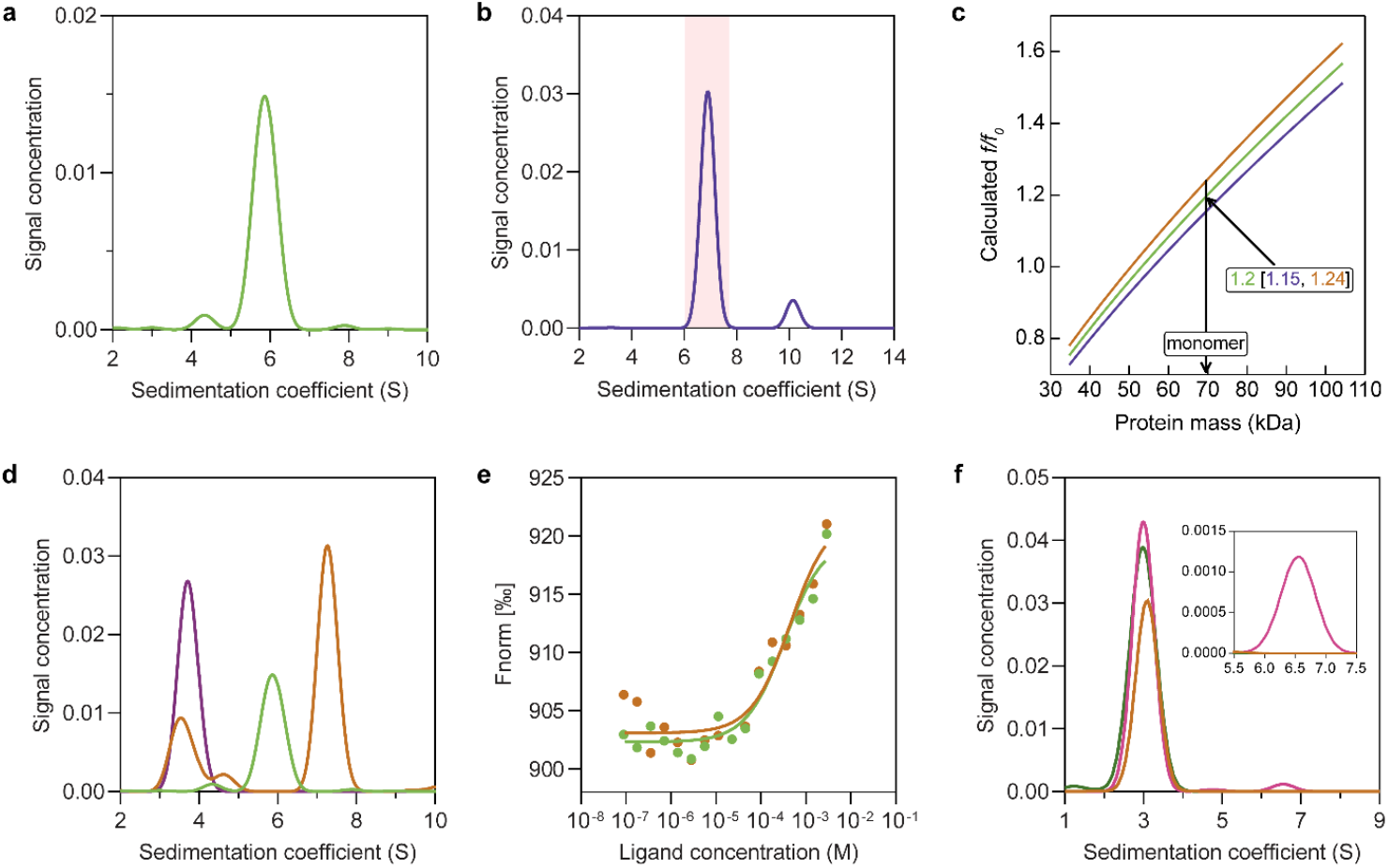
The interactions between SiaQM: SiaP and 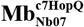, and the SiaQ: SiaM complex stoichiometry. **a**, Sedimentation velocity data for SiaQM in amphipol demonstrates a dominant peak at a sedimentation coefficient of ∼5.9 S, consistent with a single species in solution. **b**, Sedimentation velocity data for SiaQM in L-MNG (0.002%) shows a dominant monomeric species. A small proportion appears to exist as a dimer of heterodimers—which may be due to micelle cohabitation. **c**, Sedimentation data analysis^56^ demonstrates that the major species at 6.5 S in **b** (shaded) is most consistent with a monomeric transport unit of SiaQM (that is, one SiaQ and one SiaM form the complex). After determining the amount of L-MNG bound to the protein with laser interferometry, the calculated *f*/*f*_0_ for a monomer (heterodimer) for the major species in **b** is 1.2 (1σ error = 1.15−1.24), consistent with a protein in a detergent micelle. The calculated *f*/*f*_0_ for a dimer (heterotetramer) for the major species in **b** is 1.9 (1σ error = 1.83−1.97) (not shown), which is much less likely. The calculated mass of the protein from **b** was 68.2 kDa, consistent with a monomeric mass of 69.5 kDa determined from the protein sequence. The protein-detergent complex was calculated to have 83 ± 4 molecules of L-MNG bound for a total mass of the sedimenting complex of ∼152 kDa. **d**, Sedimentation velocity data for SiaQM in amphipol (green), 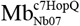 (purple) and combined (1:1.5 ratio, orange) shows a distinct increase in sedimentation coefficient for the bound species. The peak at ∼3.5 S is excess unbound megabody. **e**, Microscale thermophoresis data for the interaction between SiaQM and SiaP in the presence of Neu5Ac. Unlabelled SiaP was titrated (88.5 nM to 2.9 mM) against 20 nM RED-NHS-labelled SiaQM in amphipol. Binding saturation was not reached, indicating that saturation requires greater than 2.9 mM SiaP. Replicates fitted a dissociation constant (*K*_D_) of 341 ± 111 μM and 451 ± 206 μM. **f**, Sedimentation data for the interaction between SiaQM and SiaP in the presence of Neu5Ac. SiaQM in detergent (30 μM) incubated with 60 μM FITC-labelled SiaP (pink). The shift of ∼3% of the signal to a species at 6.5 S (inset) indicates that ∼6% SiaQM is bound to SiaP (∼1.8 μM), consistent with a *K*_D_ of ∼400 μM. Labelled SiaP without SiaQM (orange) sediments entirely at 3 S, suggesting that the species at 6.5 S is bound complex. SiaQM in amphipol (30 μM) incubated with 60 μM FITC-labelled SiaP (green). No binding was seen at these concentrations in amphipol, suggesting a high *K*_D_.

**Extended Data Figure 2.**
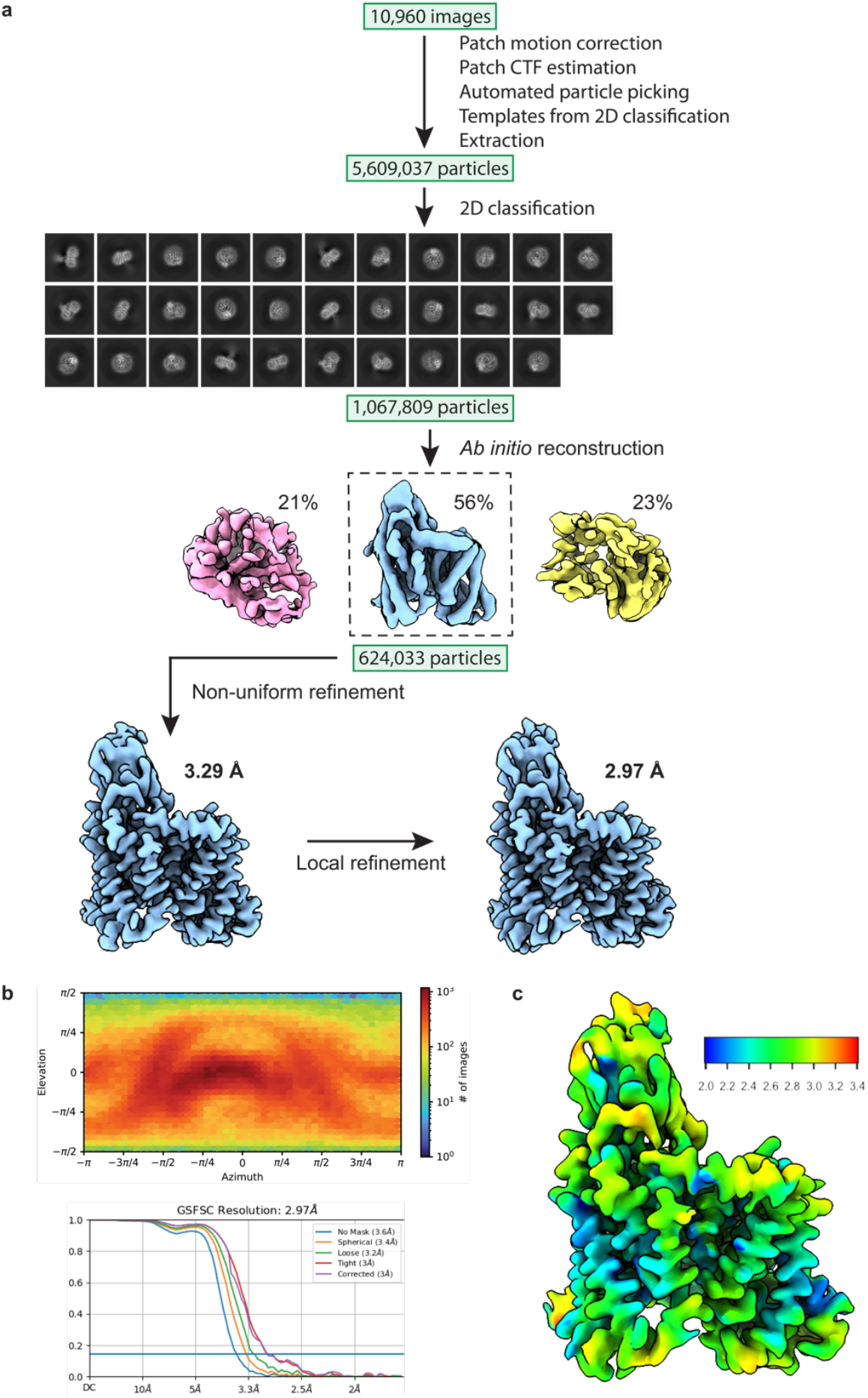
Cryo-EM workflow and analysis of the 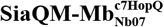complex. **a**, Workflow outlining the cryo-EM image acquisition and processing to obtain a structure of 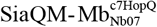 solubilised in amphipol. A representative micrograph and the selected 2D class averages used for *ab initio* reconstruction are shown. The *ab initio* reconstruction was separated into three classes to remove junk particles and the best 3D reconstruction was used as a reference model for non-uniform refinement. A mask was made in RELION 3.0^57^ and the maps were further refined with iterative rounds of local refinement. **b**, Euler angle distribution plot (top) and Fourier shell correlation (FSC) curves for the final 3D reconstruction of 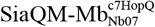. **c**, Local resolution map of 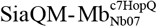 showing that the data extends to 2.2 Å in some areas.

**Extended Data Figure 3.**
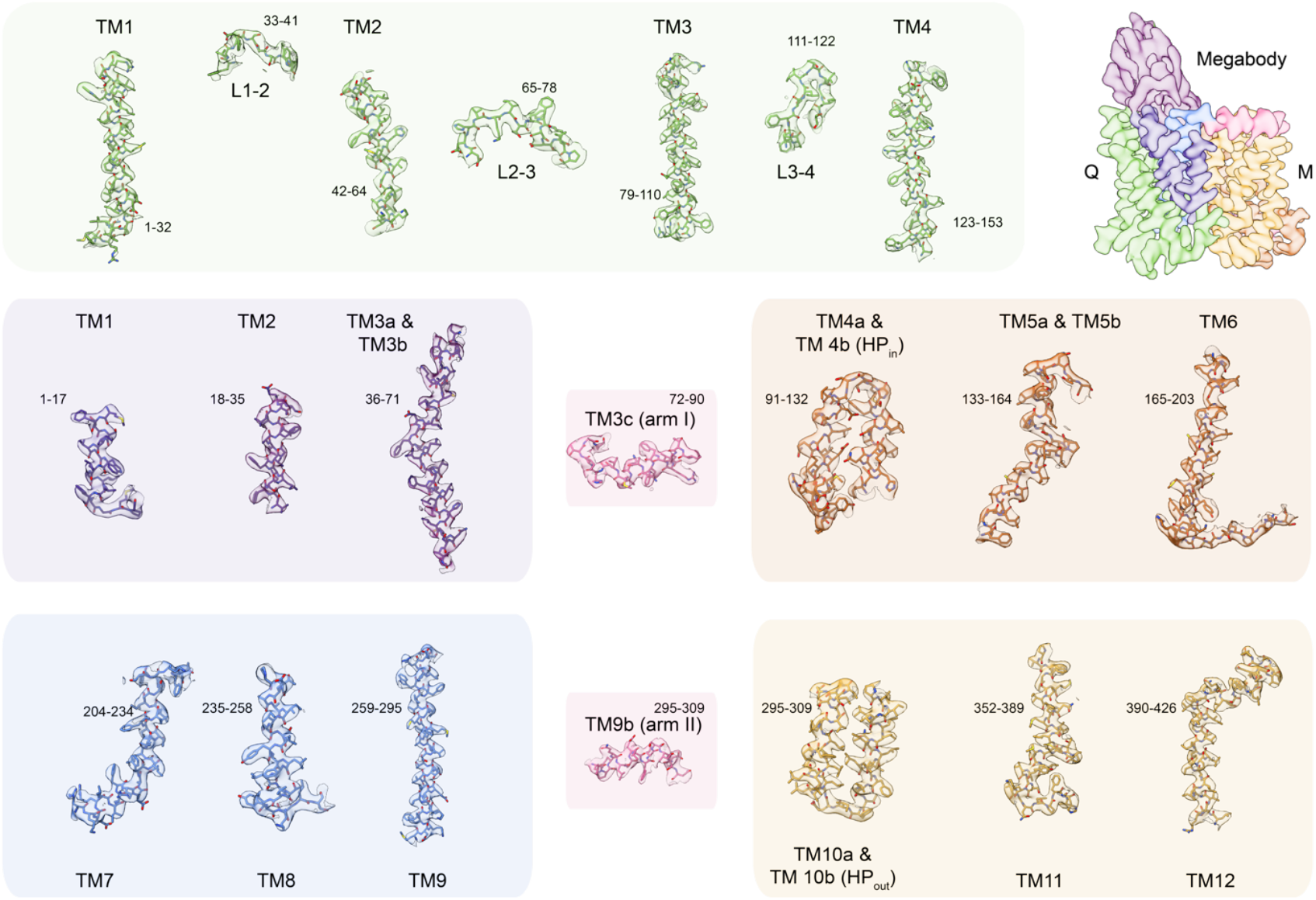
Cryo-EM density of the helices of SiaQ and SiaM. The map level is set at 7.5 σ, as calculated by *ChimeraX*^37^. The helices are coloured as in **Fig. 1b and c**.

**Extended Data Figure 4.**
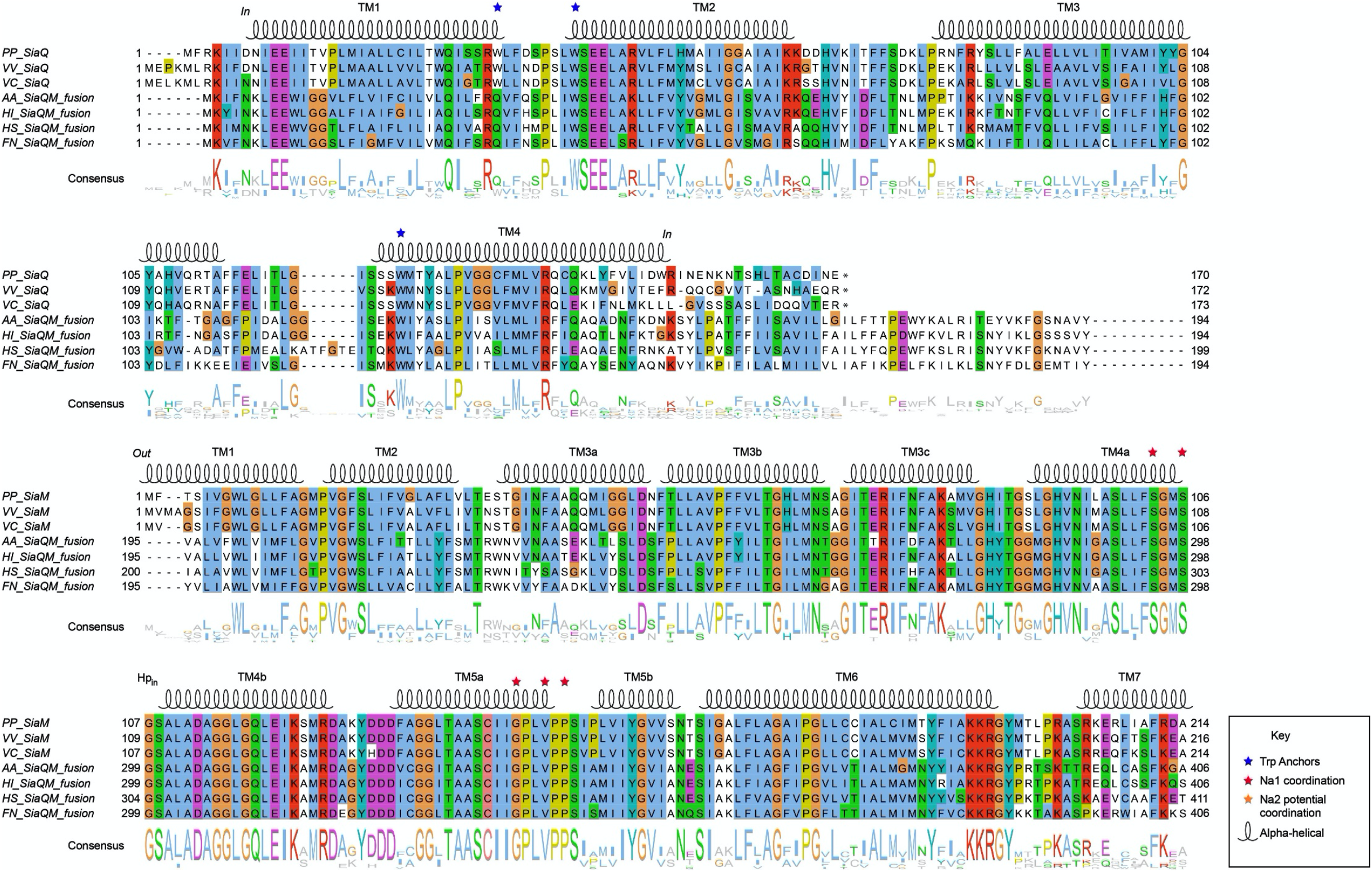
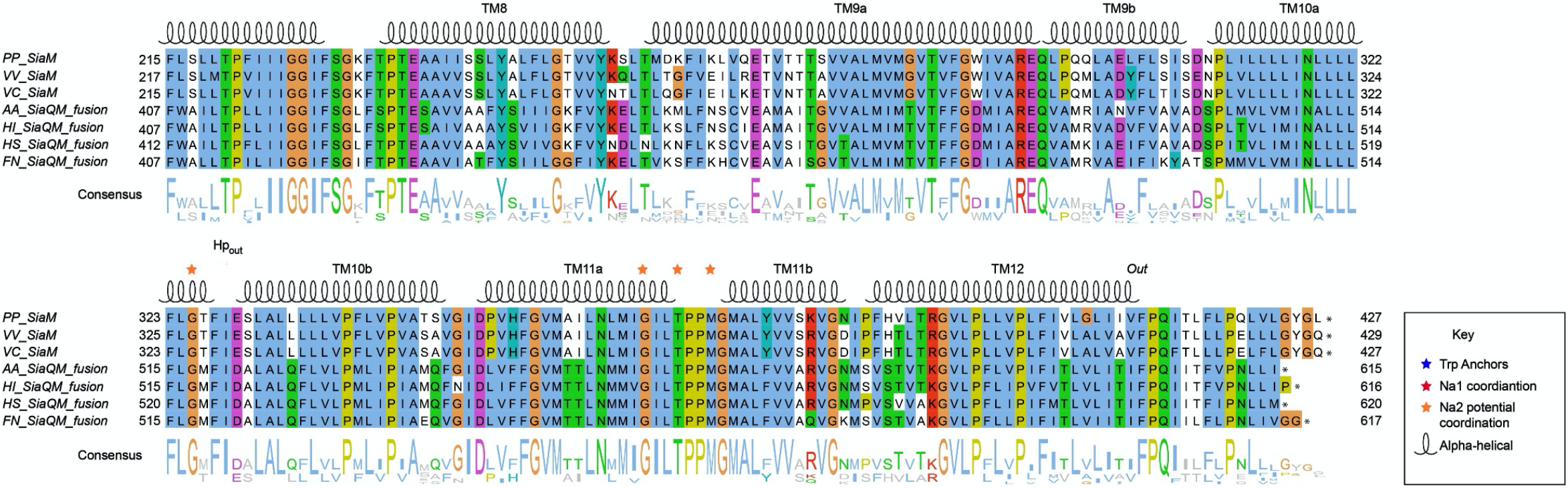
SiaQM protein sequence alignment. SiaQM protein sequences from *Photobacterium profundum (PP), Vibrio vulnificus (VV), Vibrio cholerae (VC), Aggregatibacter actinomycetemcomitans (AA), Haemophilus influenzae (HI), Histophilus somni (HS)* and *Fusobacterium nucleatum (FN)*. Protein sequences were aligned using TM-align. Aligned sequences were manually adjusted where appropriate and coloured in *Jalview* with *clustalx* colouring. Blue stars indicate anchoring tryptophan residues, red stars indicate Na1 coordinating residues and orange stars indicate the speculated Na2 coordinating residues.

**Extended Data Figure 5.**
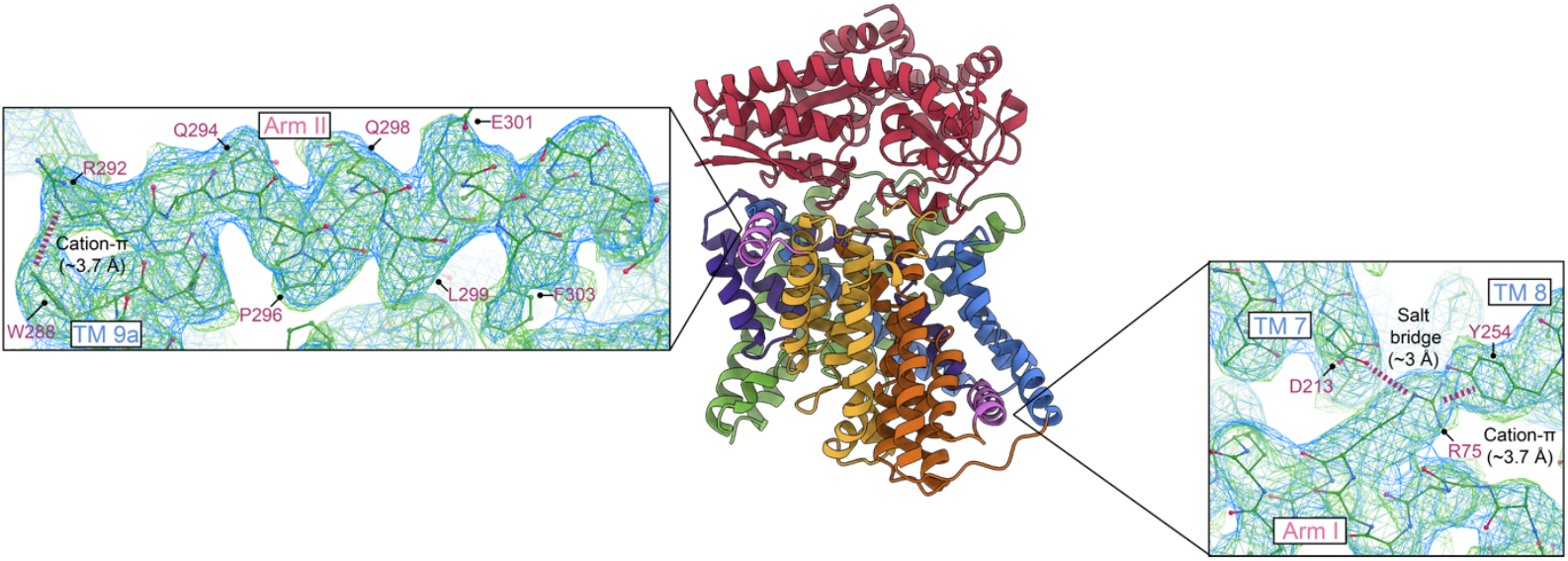
Interactions between the arm and scaffold helices. Left, inset, a cation-π interaction (W288–R292) is evident at the junction of TM9a (scaffold II) and the amphipathic arm helix II. Labelled residues show the amphipathic nature of the arm helix. Right, inset, the highly conserved R75 has a cation-π interaction with Y254 on TM8 (scaffold II), and an electrostatic interaction with D213 on TM7 (scaffold II). Cryo-EM density is shown in blue, contoured at 5σ and a locally sharpened map generated by *phenix*^46^ is shown in green, contoured at 6σ.

**Extended Data Figure 6.**
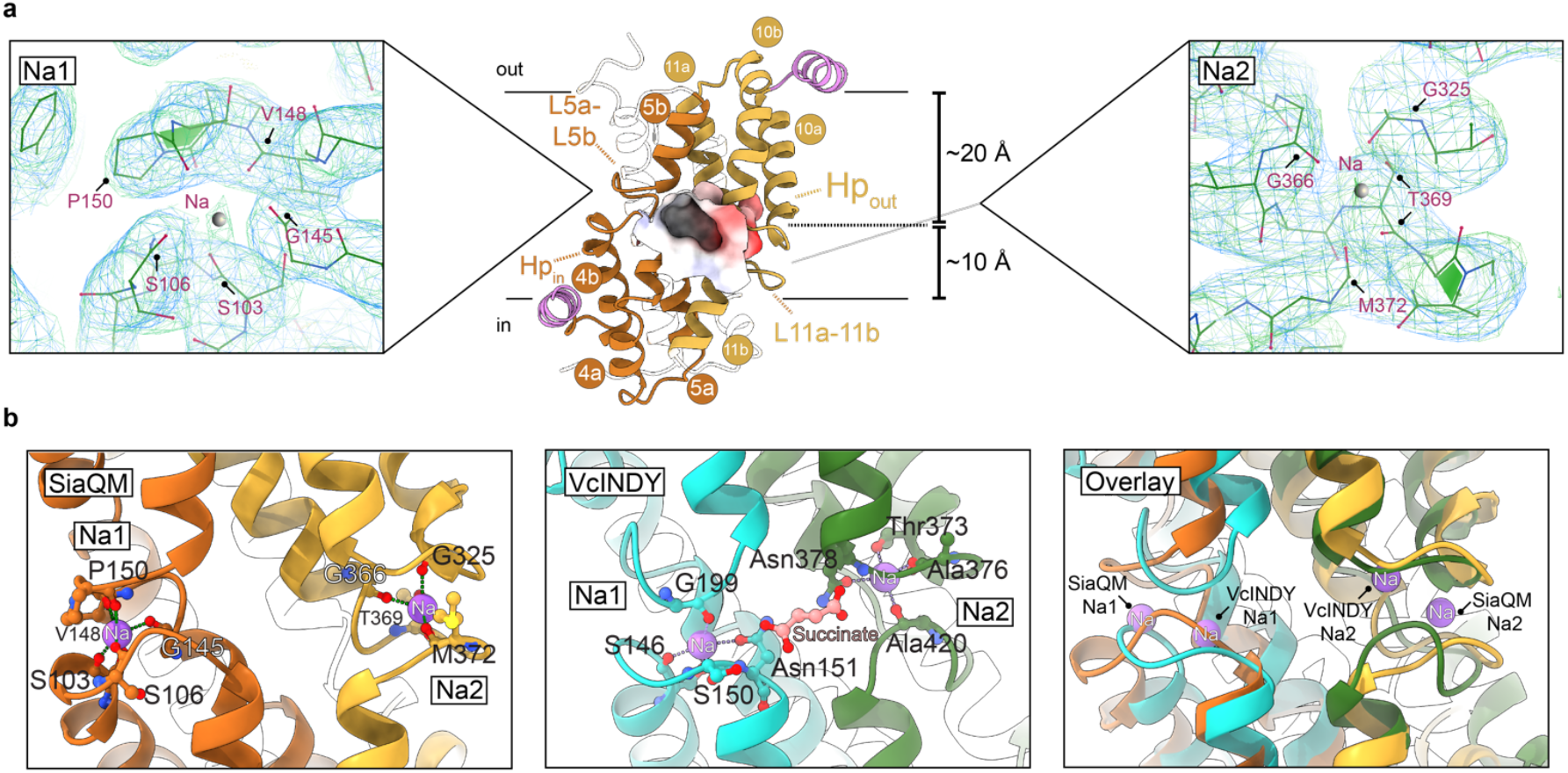
Cutaway showing the substrate-binding cavity and sodium sites. **a**, The depth of the substrate-binding site is indicated, as estimated from membrane thickness calculations^58^. The cavity (*middle*), coloured by electrostatic surface, is flanked by the predicted sodium sites, Na1 and Na2. The Na1 site is shown inset (*left*), with the cryo-EM density in blue, contoured at 5σ and a locally sharpened map generated by *PHENIX*^46^ shown in green, contoured at 6σ. The Na2 site is shown inset (*right*), with potential coordinating residues indicated. At both sites are conserved twin proline motifs, modelled here in the *cis* conformation (supported by Alphafold2 predictions). We note that the deposited model does not contain sodium ions as the deposited map without local sharpening does not show supporting density at these sites. **b** The assignment of the sodium sites (*left*) with likely coordinating residues is supported by structural comparison with VcINDY (PDB id: 5ul7) (*middle, right*).

**Extended Data Figure 7.**
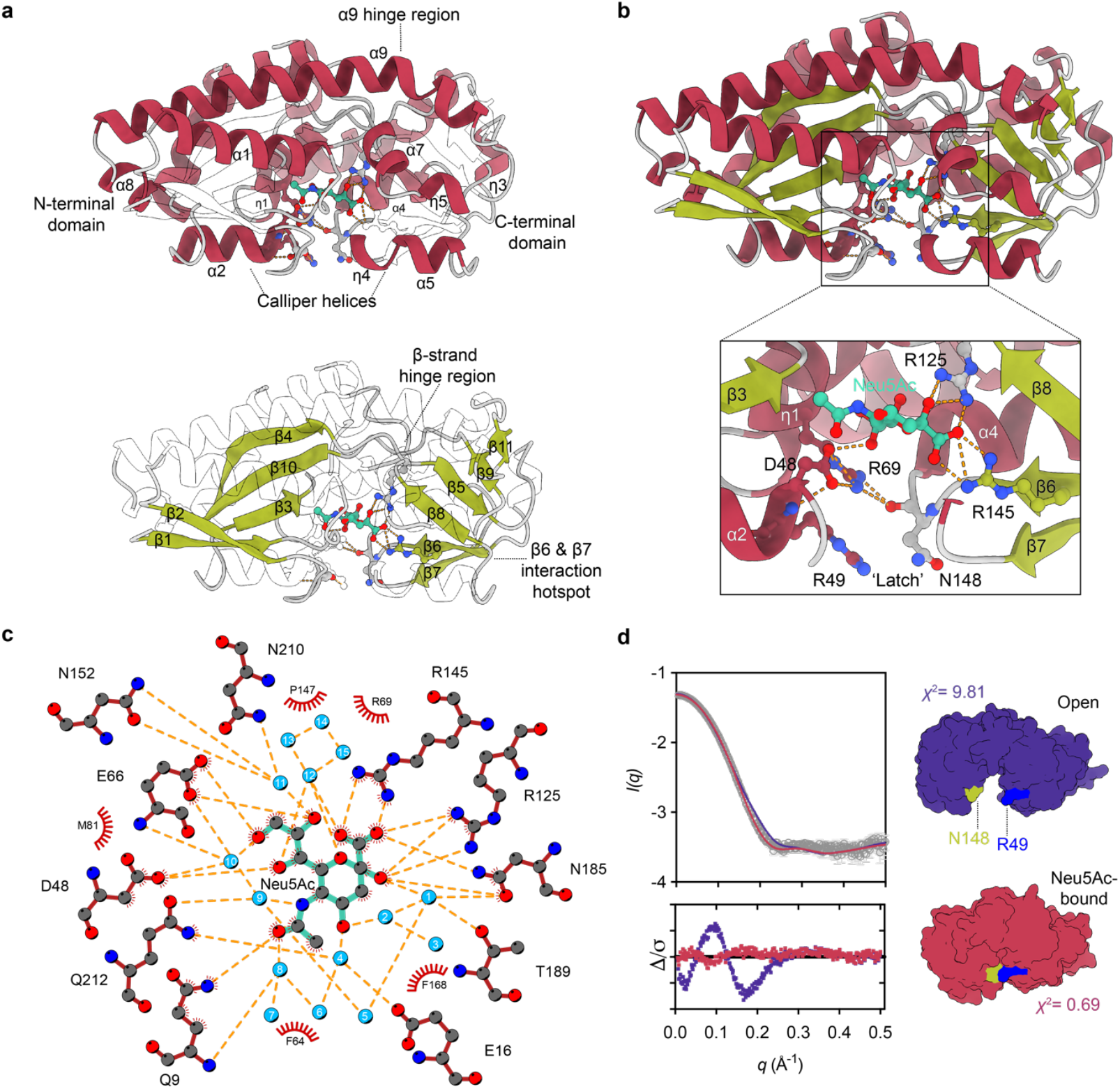
Structure of SiaP. **a**, Structural features of SiaP, including the ‘mixed-hinge’ comprised of a kinked helix α9 and two twisted β-strands (one comprised β4 and β5, and the other β9 and β10). **b**, Hydrogen bonding networks from a surface ‘latch’ area to Neu5Ac (cyan) in the binding cleft. D48 appears to play a central role, interacting with residues from both N- and C-terminal domains, and also with the hydroxyl of C7 in Neu5Ac, and a highly ordered water (10, in Ligplot). **c**, Ligplot^59^ showing close interactions at the SiaP substrate-binding site. Waters are arbitrarily numbered 1-15. **d**, SiaP small angle X-ray scattering data, showing that the X-ray crystal structure of SiaP-Neu5Ac closely resembles the conformations of bound SiaP in solution. In purple is a homology model of SiaP in an open state generated with *Modeller*^60^, using PDB ID:4mag as a template, and in maroon, the SiaP closed structure reported here. In green and blue are ‘latch’ residues, illustrating their proximity in the closed structure.

**Extended Data Figure 8.**
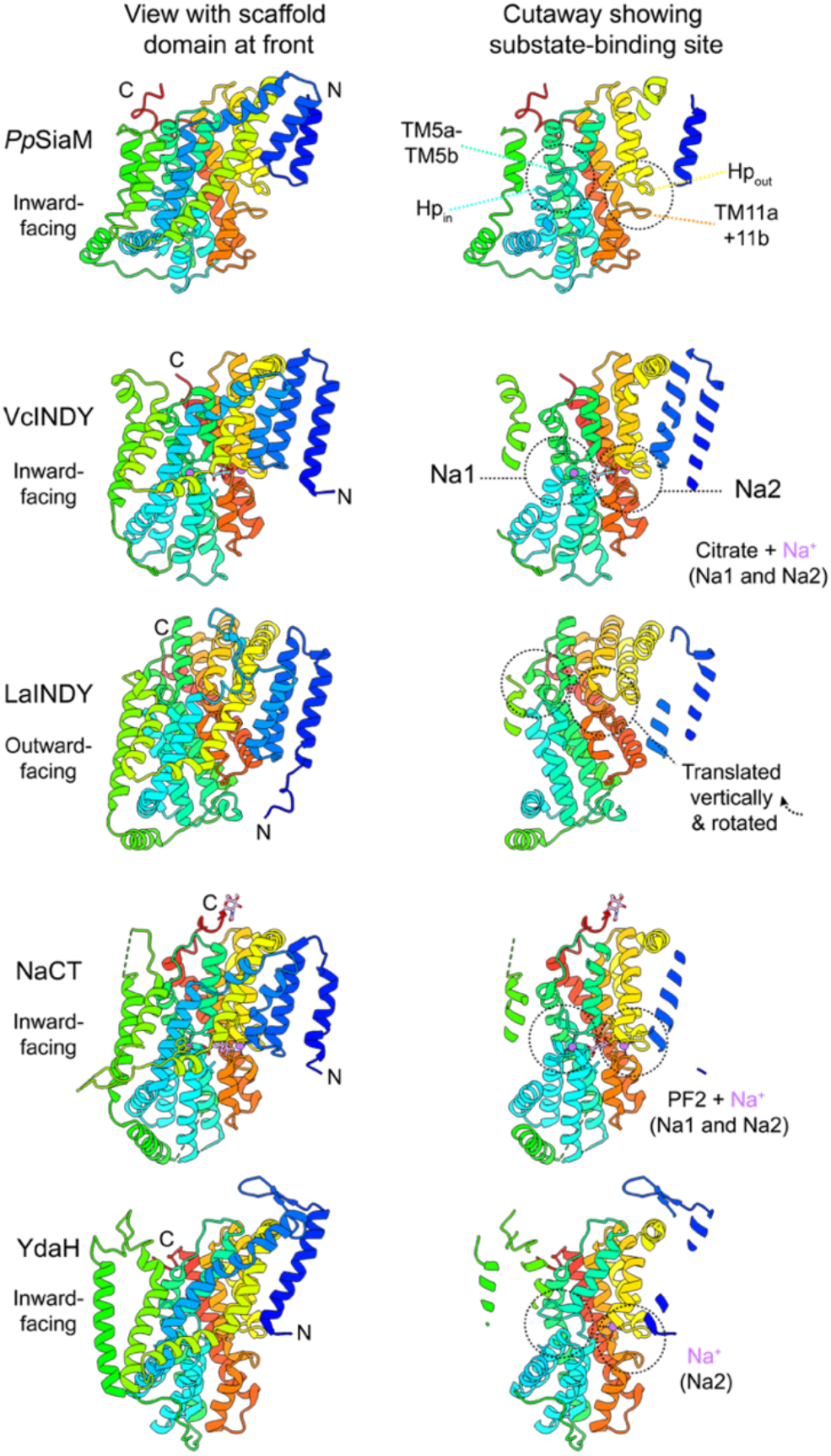
Comparison of the M-subunit fold with other elevator transporters. (*Left*) Elevator transporters oriented with the scaffold region facing the reader. (*Right*) Cutaway of the same view revealing the transport domain. The two ‘clam-shell’ motifs^27^ of the transport domain are shown in dashed circles, and contribute to the substrate/Na^+^ binding site in each transporter. Each clam-shell motif is formed at the interface of a helical hairpin and the break in a discontinuous helix. In SiaM, the two clam-shells are formed by Hp_out_ and the unwound region of TM11, as well as Hp_in_, and the unwound region of TM5. VcINDY (PDB ID: 5ul9)^27^ and LaINDY (PDB ID: 6wu1)^35^ are both members of the DASS family, and belong to the wider Ion Transporter (IT) superfamily. As illustrated, LaINDY is the only structure of this fold in an outward-facing state, clearly showing an elevator-type movement in comparison with the homologous VcINDY. YdaH (PDB ID:4r0c) is a member of the AbgT family^61^, and is also a member of the wider IT superfamily. A distinguishing feature of the M-subunit of SiaQM is its topology—the other transporters displayed here all have their N-terminal helix (blue), inserted from the cytoplasm. The M-subunit shares a conserved fold with VcINDY (4.8 Å r.m.s.d. over 352 residues, PDB id: 5ul7) and NaCT (4.2 Å r.m.s.d. over 192 residues, PDB id: 7jsj), as well as the bacterial AbgT-type transporter, YdaH (4.7 Å r.m.s.d. over 216 residues, PDB id: 4r0c), which are all members of the Ion Transporter (IT) superfamily^34^. Furthermore, these transporters form homodimers at their scaffold domains (blue, yellow and green), which is where the Q-subunit binds. The displayed transporters also show other variations such as loop length, sequence insertions, and the angle and position of TMs.

**Extended Data Figure 9.**
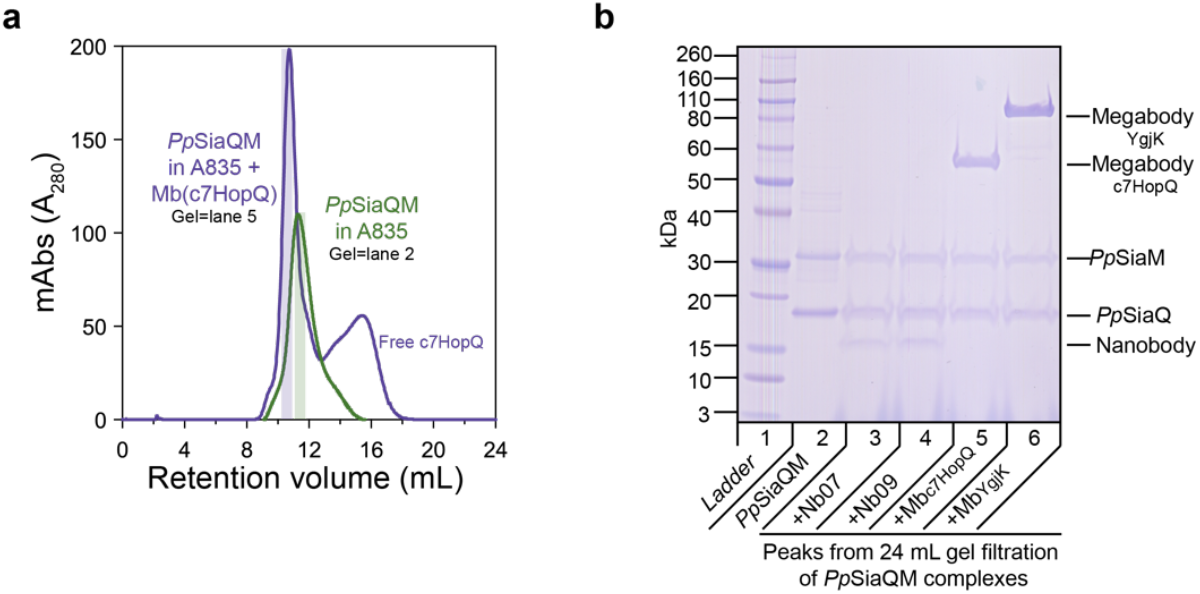
Purification of 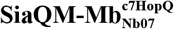 in amphipol A835 for single particle analysis. **a**, Gel filtration of SiaQM solubilised in A835 (green) and 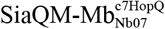 complex (purple). **b**, SDS-PAGE of peaks from size-exclusion. Lane 2 corresponds to the middle of the green peak, while Lane 5 corresponds to the middle of the purple peak, as indicated. SiaM (∼45.4 kDa) does not run true to size on SDS-PAGE, which has been documented previously^24^. An additional nanobody (Nb09, Lane 4) and megabody (MbYgjK, Lane 6) were screened but not used in this study.

**Extended Data Table 1.** Cryo-EM data collection, refinement and validation statistics.

|  | SiaQM-Mb<sup>c7HopQ</sup><sub>Nb07</sub> |
| --- | --- |
| **Data collection and processing** |  |
| Magnification | 130,000 |
| Voltage (kV) | 300 |
| Electron exposure (e<sup>-</sup>/Å<sup>2</sup>) | 71.3 |
| Defocus range (μm) | -2.0 to -0.4 |
| Pixel size (Å) | 0.6645 |
| Symmetry imposed | C1 |
| Initial particle images (no.) | 7,173,112 |
| Final particle images (no.) | 624,033 |
| Map resolution (Å) | 2.97 |
| FSC threshold | 0.143 |
| Map resolution range (Å) | 2.2 to 3.3 |
| **Refinement** |  |
| Initial model used (PDB code) | n/a |
| Model resolution (Å) | 3.0 |
| FSC threshold | 0.143 |
| Model resolution range (Å) | 2.86–3.31 |
| Model composition |  |
| Non-hydrogen atoms | 5280 |
| Protein residues | 684 |
| *B* factors (Å<sup>2</sup>) |  |
| Protein | 143.3 |
| R.m.s. deviations |  |
| Bond lengths (Å) | 0.003 |
| Bond angles (°) | 0.595 |
| Validation |  |
| MolProbity score | 1.49 |
| Clashscore | 6.68 |
| CaBLAM outliers (%) | 1.04 |
| Poor rotamers (%) | 0 |
| EMRinger score | 2.37 |
| Ramachandran plot |  |
| Favoured (%) | 97.35 |
| Allowed (%) | 2.65 |
| Disallowed (%) | 0 |
| PDB | 7qha |
| EMDB | EMD-13968 |

**Extended Data Table 2.**
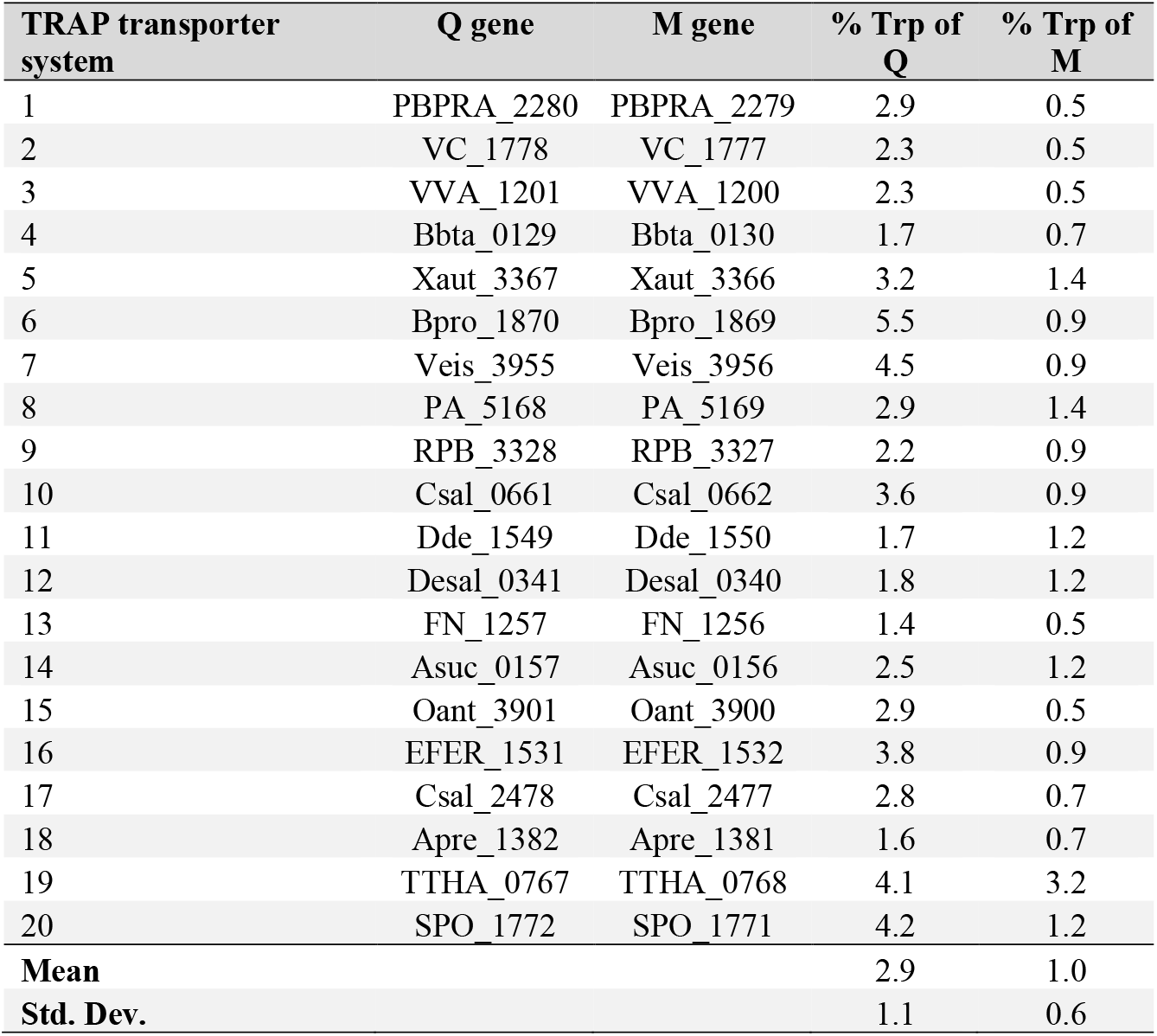
Enrichment of tryptophan residues in TRAP transporter sequences.

**Extended Data Table 3.** Data collection and refinement statistics (molecular replacement).

|  | SiaP-Neu5Ac* |
| --- | --- |
| **Data collection** |  |
| Space group | $P2_12_12_1$ |
| Cell dimensions |  |
| $a, b, c$ (Å) | 58.6, 87.3, 129.4 |
| $\alpha, \beta, \gamma$ (°) | 90, 90, 90 |
| Resolution (Å) | 43.66-1.04 (1.077-1.04) |
| $R_{\text{merge}}$ | 0.073 (1.153) |
| $I / \sigma I$ | 9.8 (1.3) |
| Completeness (%) | 99.6 (93.0) |
| Redundancy | 6.5 (5.7) |
| **Refinement** |  |
| Resolution (Å) | 1.04 |
| No. reflections | 316,707 (31,310) |
| $R_{\text{work}} / R_{\text{free}}$ | 0.1237 / 0.1386 |
| No. atoms |  |
| Protein | 5379 |
| Ligand/ion | 88 |
| Water | 799 |
| $B$-factors (Å<sup>2</sup>) | |
| Protein | 12.7 |
| Ligand/ion | 12.5 |
| Water | 26.4 |
| R.m.s. deviations |  |
| Bond lengths (Å) | 0.009 |
| Bond angles (°) | 1.08 |
| PDB | 7t3e |

2 The data was collected from a single crystal. \*Values in parentheses are for highest-resolution shell.

**Extended Data Table 4.** Interfacial/coevolved residue predictions for SiaQ:SiaM.

| Prediction method* | Rank<sup>†</sup> | Predicted probability | Residue pairs |  | Cβ-Cβ distance<sup>‡</sup> in structure (Å) |
| --- | --- | --- | --- | --- | --- |
|  |  |  | SiaM | SiaQ |  |
| *RaptorX complex-contact* | 1 | 0.56 | Glu-237 | Ile-29 | 5.5 |
|  | 2 | 0.54 | Ile-240 | Leu-25 | 4.7 |
|  | 3 | 0.54 | Ile-240 | Ile-29 | 6.3 |
|  | 4 | 0.54 | Lys-232 | Trp-33 | 8.1 |
|  | 5 | 0.54 | Ala-42 | Tyr-103 | 4.4 |
|  | 30 | 0.49 | Gln-267 | Val-15 | 6.8 |
|  | 31 | 0.49 | Gly-15 | Lys-69 | 5.4 |
|  | 32 | 0.49 | Ser-243 | Leu-22 | 7.3 |
|  | 33 | 0.49 | Val-18 | Val-68 | 5.8 |
|  | 34 | 0.49 | Val-270 | Met-53 | 6.8 |
| *GREMLIN* | 1 | 1 | Leu-244 | Thr-26 | 6.3 |
|  | 2 | 1 | Ile-23 | Thr-96 | 5.6 |
|  | 3 | 1 | Asp-50 | Arg-47 | 6.4 |
|  | 4 | 1 | Lys-232 | Trp-33 | 8.1 |
|  | 5 | 1 | Thr-271 | Lys-64 | 6.8 |
|  | 30 | 0.91 | Met-45 | Met-100 | 7.9 |
|  | 31 | 0.9 | Leu-49 | Leu-51 | 5.5 |
|  | 32 | 0.89 | Ile-240 | Thr-26 | 6.6 |
|  | 33 | 0.89 | Ala-277 | Phe-50 | 8.8 |
|  | 34 | 0.89 | Met-45 | Ile-55 | 9.8 |

\* Each prediction method was run with default settings.
<sup>†</sup> The residue pairs are ranked according to each method's estimate of the probability that a residue pair is in contact. The top five residue pairs are displayed, alongside the residue pairs ranking 30-34 with a lower predicted coupling probability, arbitrarily chosen to show the extent of useful information provided by these analyses. We note that the probabilities are calculated differently between methods, though we have chosen to display them here so they can be compared to the interfacial predictions for SiaM:SiaP, below.
<sup>‡</sup> Cβ-Cβ distance was analysed (Cα-Cβ for residue pairs containing glycine) as residue pairs are typically defined to be in contact if their Cβ-Cβ distance is less than 8 Å.

**Extended Data Table 5.** Interfacial/coevolved residue predictions for SiaM:SiaP and SiaQ:SiaP.

| Prediction method* | Rank<sup>†</sup> | Predicted probability | Residue pairs |  | Residue location (SiaM) | Residue location (SiaP) | Location colour (in Fig. 3a) |
| --- | --- | --- | --- | --- | --- | --- | --- |
|  |  |  | SiaM | SiaP |  |  |  |
| *RaptorX complex-contact* | 1 | 0.49 | Ser-164 | Glu-197 | L5b-6 | η5 | Orange (A) |
|  | 2 | 0.49 | Thr-234 | Ala-179 | L7-8 | Lα5-β7 | Blue (C) |
|  | 3 | 0.49 | Pro-235 | Leu-173 | Tm8 | α5 | Blue (C) |
|  | 4 | 0.48 | Thr-234 | Ala-174 | L7-8 | α5 | Blue (C) |
|  | 5 | 0.48 | Ala-291 | Gln-44 | Tm9a | Lβ2-α2 | Blue (C) |
| *GREMLIN* | 1 | 1 | Gly-166 | Tyr-172 | Tm6 | α5 | Orange (A) |
|  | 2 | 1 | Gln-294 | Leu-38 | L9a-9b | β2 | Blue (B) |
|  | 3 | 0.97 | Ser-164 | Glu-197 | L5b-6 | η5 | Orange (A) |
|  | 4 | 0.97 | Gly-47 | Ala-50 | Tm3a | α2 | Purple (D) |
|  | 5 | 0.86 | Pro-235 | Leu-173 | Tm8 | α5 | Blue (C) |
| Prediction method* | Rank<sup>†</sup> | Predicted probability | Residue pairs |  | Residue location (SiaQ) | Residue location (SiaP) | Location colour (in Fig. 3a) |
|  |  |  | SiaQ | SiaP |  |  |  |
| *RaptorX complex-contact* | 1 | 0.49 | Thr-118 | Pro-165 | L3-4 | β6 | Green (E) |
|  | 2 | 0.49 | Ile-117 | Pro-165 | L3-4 | β6 | Green (E) |
|  | 3 | 0.48 | Pro-38 | Ser-162 | L1-2 | β6 | Green (F) |
|  | 4 | 0.47 | Thr-118 | Ala-149 | L3-4 | Lβ5-α4 | Green (E) |
|  | 5 | 0.45 | Leu-40 | Pro-163 | L1-2 | β6 | Green (F) |
| *GREMLIN* | 1 | 1 | Pro-38 | Ser-162 | L1-2 | β6 | Green (F) |
|  | 2 | 0.63 | Arg-110 | Met-57 | Tm3 | α2-β3 | Green (E) |
|  | 3 | 0.59 | Arg-110 | Ser-56 | Tm3 | α2 | Green (E) |
|  | 4 | 0.55 | Leu-40 | Pro-163 | L1-2 | β6 | Green (F) |
|  | 5 | 0.47 | Arg-110 | Asp-219 | Tm3 | α8 | Green (E) |

**Extended Data Table 6.**
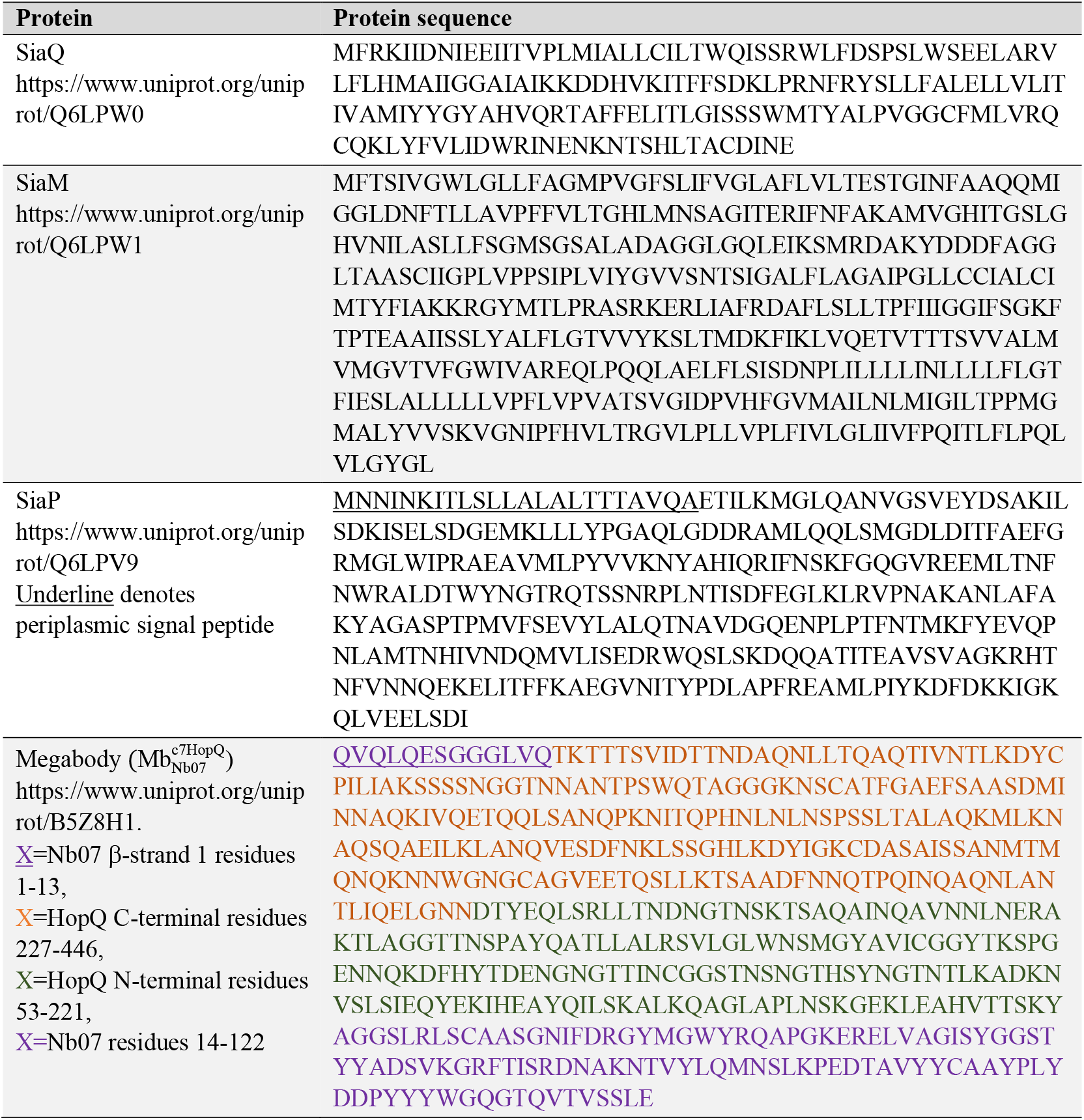
Protein sequences of SiaQM and SiaP expressed and purified in this study.

